# Distinctions among real and apparent respiratory motions in human fMRI data

**DOI:** 10.1101/601286

**Authors:** Jonathan D. Power, Benjamin M. Silver, Marc J. Dubin, Alex Martin, Rebecca M. Jones

**Author notes:** **Corresponding author:** Jonathan D Power. **Contact information:** Jonathan Power, Benjamin Silver, Marc Dubin, Alex Martin, Rebecca Jones.

## Abstract

Head motion estimates in functional magnetic resonance imaging (fMRI) scans appear qualitatively different with sub-second image sampling rates compared to the multi-second sampling rates common in the past. Whereas formerly the head appeared still for much of a scan with brief excursions from baseline, the head now appears to be in constant motion, and motion estimates often seem to divulge little information about what is happening in a scan. This constant motion has been attributed to respiratory oscillations that do not alias at faster sampling rates, and investigators are divided on the extent to which such motion is “real” motion or only “apparent” pseudomotion. Some investigators have abandoned the use of motion estimates entirely due to these considerations. Here we investigate the properties of motion in several fMRI datasets sampled at rates between 720-1160 ms, and describe 5 distinct kinds of respiratory motion: 1) constant real respiratory motion in the form of head nodding most evident in vertical position and pitch, which can be very large; 2) constant pseudomotion at the same respiratory rate as real motion, occurring only in the phase encode direction; 3) punctate real motions occurring at times of very deep breaths; 4) a low-frequency pseudomotion in only the phase encode direction following very deep breaths; 5) slow modulation of vertical and anterior-posterior head position by the respiratory envelope. We reformulate motion estimates in light of these considerations and obtain good concordance between motion estimates, physiologic records, image quality measures, and events evident in the fMRI signals.

**Highlights:**

- Examines several fast-TR datasets with sampling rates of 720-1160 ms
- Identifies 7 kinds of motion in fMRI scans, 5 of them related to respiration
- Identifies 2 forms of pseudomotion occurring only in phase encode direction
- Pseudomotion is a function of soft tissue mass, not lung volume
- Reformulates motion estimates to draw out particular kinds of motion

## Introduction

Head motion causes major signal disruptions in functional magnetic resonance imaging (fMRI) data (Caballero-Gaudes and Reynolds, 2017; Friston et al., 1996). Head motion causes modeling confounds in task fMRI when it correlates with task design (Barch et al., 1999; Bullmore et al., 1999), and it clouds interpretation of resting state fMRI studies because it adds spatially focal covariance to images (Horien et al., 2018; Satterthwaite et al., 2017). Though some motion may be voluntary, some is involuntary, arising from head nodding during respiratory cycles, head bobbing with cardiac pulsations, or events like yawns or sighs or twitches or sneezes, in addition to other factors (Zaitsev et al., 2017). In previous decades, when whole-brain fMRI images were typically obtained only every few seconds, little could be said about the extent to which various motions reflected various physiological or voluntary causes in a typical dataset. However, multi-band accelerations now permit much more rapid brain imaging, and sub-second acquisitions are becoming normative. Such rapid acquisitions are yielding head position estimates fundamentally different than prior estimates, as noted by several authors (Glasser et al., 2018; Power, 2017). Figure 1 illustrates the difference using data from the Brain Genomics Superstruct Project (GSP) and the Human Connectome Project (HCP) (Holmes et al., 2015; Van Essen et al., 2013). In older datasets like the GSP with multi-second sampling rates it was common for the head to appear essentially still for much of the scan, with occasional spikes of motion. However, in datasets like the HCP, with sub-second sampling rates, motion estimates appear much “noisier”, and in many cases the head appears to be in constant motion.

**Figure 1:**
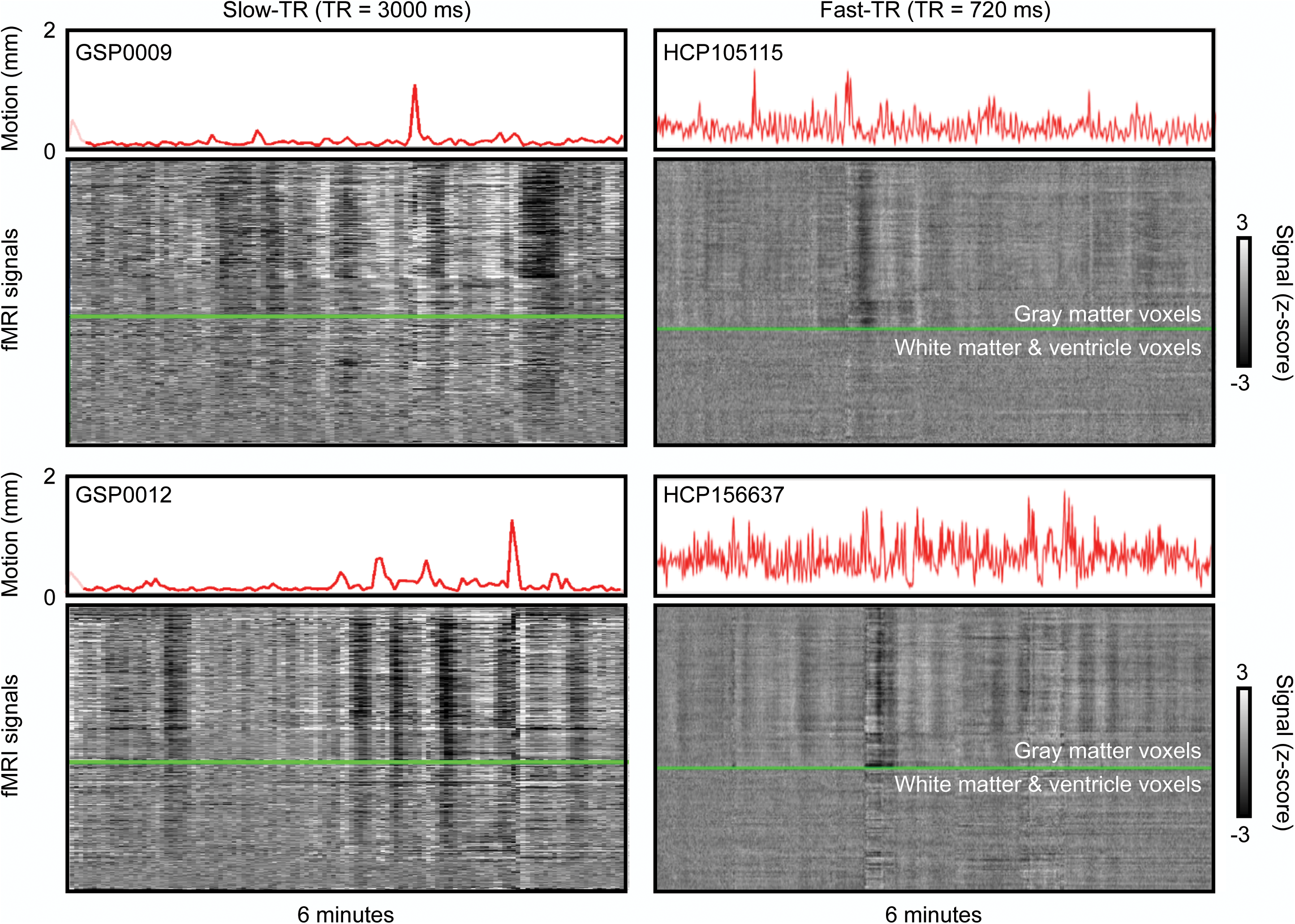
Illustrations of motion in slow-TR vs fast-TR data. Left column shows two 6-minute scans from the Brain Genomics Superstruct Project (GSP), right column shows two 6-minute segments of Human Connectome Project data. Slow-TR scans show “floors” of motion in many scans, where the brain appears essentially still, with occasional excursions from the floor. By contrast, motion is continuously elevated throughout many HCP scans. Images are modified from supplemental videos of {Power, 2017}.

Some authors have attributed this noisy appearance of head motion in fast-repetition-time (fast-TR) data to the fact that some physiological cycles no longer alias, in particular respiratory cycles (Burgess et al., 2016; Siegel et al., 2017). Breathing can cause the head to move, but changes in lung volume can also cause a particular kind of artifact called pseudomotion, which manifests as a shift of the brain when the lung expands (Brosch et al., 2002; Durand et al., 2001; Raj et al., 2001). Some authors have postulated that the HCP data, and fast-TR data generally, are substantially contaminated by respiratory pseudomotion and have proposed either filters to remove the motion (Fair et al., 2018), or to disregard motion estimates entirely and measure “motion” in terms of where denoising strategies intervene in the timeseries (Glasser et al., 2018). Because motion of the head is hardly ever measured externally, and because respiratory records are also infrequently obtained during scanning, there has been little evidence to inform the extent to which motion in fast-TR data is “real” or not. However, we recently found that physical restraint of heads by anatomy-specific molds reduced head motion amplitude in images, and evidence of motion artifact, and that this occurred even in the low-motion portions of scans where only ongoing respiratory motions were apparent (Power et al., 2019). These findings are compatible with a reduction in real motion, and are hard to explain by pseudomotion, prompting us to examine the sources of motion in fast-TR data more closely.

In this article we examine scans from several fast-TR datasets and discern five distinct kinds of respiratory motion: true constant motion of the head at the respiratory rate, false constant pseudomotion of the head at the respiratory rate, true intermittent spikes in motion due to sighs and yawns, a distinct and very slow form of pseudomotion at the times of signs and yawns, and long-term modulation of head position by respiratory depth. We reformulate motion estimates for fast-TR data, and demonstrate how reformulation reveals distinct kinds of relationships between motion and respiration and body habitus.

## Methods

### Data

The “900-subject” release of the Human Connectome Project data was obtained, with a focus on the following files in each subject:

Four resting state fMRI scans transformed to atlas space (in each subject’s /MNINonLinear/Results folder): [RUN]=REST1_LR, REST1_RL, REST2_LR, REST2_RL (this order is runs 1-4 in the text). rfMRI_[RUN].nii.gz and rfMRI_[RUN]_hp2000_clean.nii.gz scans were obtained, representing minimally preprocessed and FIX-ICA-denoised data. For each of these scans, the [RUN]_Physio_log.txt and Movement_Regressors_dt.txt files were also obtained.

Structural scans transformed to atlas space (in each subject’s /MNINonLinear/ folder): the T1w.nii.gz and the aparc+aseg.nii.gz files, representing the anatomical T1-weighted scan and its FreeSurfer segmentation.

### Image Processing

The aparc+aseg.nii.gz file for each subject underwent a set of serial erosions within white matter and ventricle segments, exactly as in (Power et al., 2017b). Masks of cortical gray matter, the cerebellum, and subcortical nuclei were extracted, as were serially eroded layers of superficial, deep, and deepest (with respect to distance from gray matter) masks of the white matter and ventricles. These masks, together, include all in-brain voxels of these tissue types, and are used to extract certain signals and to order signals for “gray plots” (Power, 2017).

For the purpose of making useful gray plots, because of the considerable thermal noise in HCP scans, a within-mask 6 mm FWHM Gaussian kernel was applied to the data using the above masks (illustrated for HCP data in (Power, 2017)). This blurring does not mix tissue compartments due to the use of masks.

### Parameter processing

#### Respiratory and cardiac measures

Respiratory belt and pulse oximeter traces (sampled at 400 Hz) first underwent visual inspection in their entirety to determine if the quality was sufficient for reliable peak detection, since traces are often partially or fully corrupted. Only subjects with traces deemed likely to successfully undergo peak detection in all runs were analyzed (440 of the original 900 passed this quality control step; traces and quality decisions can be seen in the materials of (Power, 2019)^1^.).

After selection, for respiratory traces, an outlier replacement filter was used to eliminate spurious spike artifacts (Matlab command: filloutliers(resp_trace,‘linear’,‘movmedian’,100)) and the traces were then gently blurred to aid peak detection (Matlab command: smoothdata(resp_trace,‘sgolay’,400)) (a 1-second window for a 400 Hz signal). These treated respiratory traces are the ones shown in Figures unless indicated.

Following prior literature, several respiratory measures were derived from the treated respiratory belt trace. First, the (upper) envelope of the trace over a 10-second window (at 400 Hz) was calculated after (Power et al., 2018) (Matlab command: envelope(zscore_resp_trace,4000,‘rms’)). Second, the RV measure, defined as the standard deviation of the treated respiratory trace within a 6-second window, was calculated following (Chang and Glover, 2009) (Matlab command: movstd(zscore_resp_trace, 2400,‘endpoints’,‘shrink’)). Finally, the RVT measure, defined for all peaks as ((peak-prior trough)/(time between peaks)), was calculated after (Birn et al., 2006). Peak detection on the trace yielded peaks (and troughs, using the inverted trace) for calculations (Matlab command: (findpeaks(zscore_resp_trace,‘minpeakdistance’,800,‘minpeakprominence’,.5)). The minimum peak distance presumes breaths occur more than 2 seconds apart. If a peak did not have a preceding trough prior to the previous peak, no value was scored at that peak. All traces and derived measures were visually checked to ensure that outliers and abnormalities would not drive results. These 3 measures were termed ENV, RV, and RVT in figures.

#### Data quality measures

The data quality measure DVARS was calculated after (Power et al., 2012; Smyser et al., 2010) as the root mean squared value in the brain at each timepoint of all voxel timeseries differentiated in time by backwards differences. DVARS by convention is 0 at the first timepoint.

#### Head position and head motion measures

Head position was taken from the Movement_Regressors_dt.txt files. In Figures these position parameters are displayed either after subtracting the first timepoint value from the timeseries (so that all traces start at zero) or by spreading them on the y-axis by adding constants to each trace (so that each individual trace is seen clearly). Head motion was represented by Framewise Displacement (FD) measures, following (Power et al., 2012), wherein all position measures were differentiated in time by backwards differences, rotational measures were converted to arc displacement at 5 cm radius, and the sum of the absolute value of these measures was calculated. FD is typically calculated by backwards differences to the preceding timepoint (for HCP data, 720 ms prior), but historically FD measures using sampling rates of 2-4 seconds were common; for comparison to such measures, FD was also calculated by backwards differences over 4 timepoints (e.g., 4 * 720 ms = 2.88 seconds effective sampling rate) where indicated.

#### Frequency content

At times, the frequency content of certain signals is illustrated, such as for respiratory belt traces or for position estimates. Power spectral density estimates were generated via Welch’s method (e.g., for a respiratory trace, Matlab command: [pw pf] = pwelch(signal,[],[],[],400,‘power’)) and are shown on a semilogarithmic scale. All spectra were visually checked to ensure correct peak frequency identification.

At times and where indicated, position estimates were filtered to exclude certain frequencies using a Butterworth filter (Matlab commands for an example stopband of 0.2-0.5 Hz: TR = 0.72; nyq = (1/TR)/2; stopband=[0.2 0.5]; Wn = stopband/nyq; filtN = 10; [B,A] = butter(filtN,Wn,‘stop’); filtposition(:,column) = filtfilt(B,A,position(:,column));).

### Accessory data

To examine the generalizability of certain findings, head position estimates from other fast-TR datasets were examined.

MyConnectome dataset: The data studied were motion estimates from 84 scans of a single adult scanned at rest, with TR = 1160 ms and scan durations of 10 minutes, the exact data used in (Laumann et al., 2015; Power, 2017; Power et al., 2017b). These position estimates were obtained using the 4dfp tools of Washington University, which yield similar estimates to other software packages (Power et al., 2017a). A Siemens 3T Skyra scanner with 32-channel head coil was used, and the sequence was a multi-band factor 4 EPI sequence with TR = 1160 ms, TE = 30 ms, flip angle = 63 degrees, voxel size 2.4 × 2.4 × 2 mm, and AP phase encoding direction. Respiratory records were not acquired with this dataset.

Cornell Dataset 1: N = 70 adult subjects scanned once at rest on a Siemens Prisma 3T scanner using a 32-channel head coil for 5.4 min at rest with AP phase encode direction, TR = 766 ms, TE = 34 ms, multi-band factor 8, and 2 mm isotropic voxel sizes. Realignment was calculated using AFNI’s 3dvolreg command. These scans have not been previously published and were collected with written informed consent from subjects under a protocol approved by the Weill Cornell Medicine Institutional Review Board. Respiratory records were not provided with this dataset.

Cornell Dataset 2: N = 11 pediatric subjects (ages 7-17, mean 13.7) scanned twice each at rest for 4.8 min in a Siemens Prisma 3T scanner with 32-channel head coil. Sequences used AP phase encode, TR = 800 ms, TE = 30 ms, multi-band factor 6, and 2.4 mm isotropic voxel sizes. Realignment was calculated using AFNI’s 3dvolreg command. These scans a subset of the data published in (Power et al., 2019), the “head mold off” scans of subjects younger 18 years old. The scans were collected with written informed consent from parents and assent of pediatric subjects under a protocol approved by the Weill Cornell Medicine Institutional Review Board.

### Estimating motion before image realignment

For 10 subjects of the HCP dataset and 10 sessions of the MyConnectome dataset, unprocessed images were obtained, and the center of mass was calculated in each volume, yielding position estimates in the X (left-right), Y (anterior-posterior), and Z (superior-inferior) axes.

## Results

The important kinds of measures used in this paper are illustrated in Figure 2 for portions of three HCP scans. At top in gray are the realignment parameters (shown on 2 mm vertical scale) and in red the framewise displacement (FD) motion estimate (also shown on a 2 mm scale). The respiratory belt trace is in blue in the third panel. Note the constant zigzags in the gray position traces matching visually the frequency seen in the blue respiratory traces. DVARS traces before and after FIX-ICA denoising are shown, as are respiratory measures derived from the respiratory belt (ENV, RV, and RVT). “DVARS dips” in FIX-ICA-denoised images have been proposed as a measure of motion that would be insensitive to pseudomotion (Glasser et al., 2018). DVARS spikes in minimally processed data indicate that signals are changing abnormally rapidly, and DVARS dips after FIX-ICA denoising indicate abnormally little change in signals at those times. The dips are caused by FIX-ICA intervening strongly at those timepoints to remove so much variance that unusually little variance remains, and the logic of proposing DVARS dips as an index of motion is that FIX-ICA would intervene so strongly mainly at times of motion. At bottom are the signals of all in-brain voxels of the minimally processed image shown as a heat map and ordered by anatomical compartment. These scans were chosen to illustrate a “best case” scenario at left, where FD and DVARS abnormalities are highly concordant, and then two “worse” scenarios at middle and right where there is less concordance between measures. In the middle and right scans, there is so much “noise” in the motion estimate that visible step changes in the gray position measures do not necessarily translate into noticeable motion spikes. Sometimes these changes in position have no DVARS spike, and sometimes they have a spike but no dip. These images encapsulate the uncertainty about what “motion” means in fast-TR data. Both constant and punctate motions are seen, but either might simply be pseudomotion, since many of the punctate motions (black arrows) are occurring at times of brief pauses and/or deep breaths visible in the respiratory traces.

**Figure 2:**
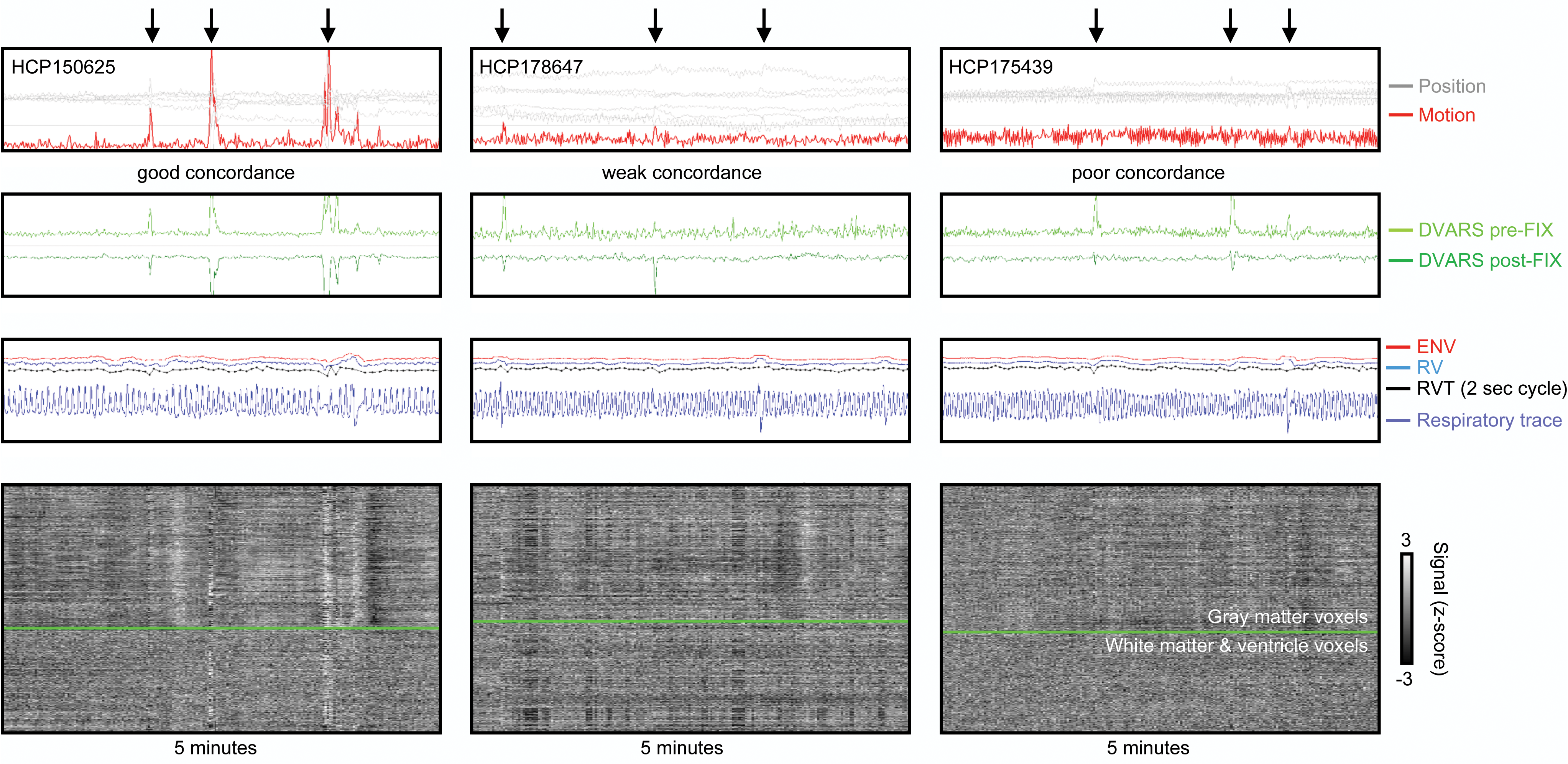
Examples of concordance and discordance of motion and DVARS. Top panels show position and motion estimates (the realignment parameters, and framewise displacement (FD), on a 2 mm vertical scale; the horizontal line is FD = 0.5 mm), the next panels show DVARS before and after FIX-ICA, the next panels show respiratory belt traces and the derived measures ENV, RV, and RVT, and bottom panels show heat maps of all in-brain voxel signals in minimally processed images. In all scans there is constant motion at the respiratory frequency in the gray position estimates and red motion estimate. In the left scan, there is good concordance between motion estimates, DVARS spikes, and DVARS dips, a “best case scenario”. However, the middle and right scans show poor concordance of motion estimates and DVARS, as well as instances of position estimates visibly changing without the changes being notable in the motion estimates (or DVARS). Black arrows mark times when position, motion, or DVARS measures suggest that motion has occurred.

### Basic properties of respiration

We will soon begin to fractionate motions based on their properties, but first it is helpful to examine the respiratory properties that are likely to manifest in the motion estimates. Breathing assumes a variety of forms within and across scans. Figure 3 illustrates respiratory traces from 8 subjects, notable for the variation in frequencies across scans and within scans. Many scans exhibit pauses in breathing, changes in the depth of breathing, occasional deep breaths, and some breathing patterns simply appear chaotic. Breathing patterns have recognizable and reproducible features within subjects. Figure 4A shows the peak respiratory frequency for each of the four scans for all 440 subjects studied in this paper. Peak frequencies almost entirely fall between 0.2-0.4 Hz, with an average of 0.3 Hz. Within-subject variance in peak frequency is 0.018 Hz, much less than the variance across the group of 0.055 Hz. A minority of subjects do display quite variable peak frequency across scans (the instances with purple stars in Figure 4A are shown in Figure S1). Some subjects have very broad spectral peaks reflecting a wide variety of respiratory rates within a run. Harmonics are visible in the spectra (i.e., the secondary peaks at twice the fundamental frequency in the 2^nd^-4^th^ panels of Figure 4B). In the examples shown, there is obvious similarity of the spectra within individuals, an observation borne out statistically in the entire dataset: correlations of the spectra (spanning 0.05-0.6Hz) are r = 0.79 ± 0.15 within subjects, versus r = 0.44 ± 0.30 across all scans (p = 0). Individual subjects sometimes exhibit characteristic events in multiple scans. For example, the top subject of Figure 4B has notable accelerations and decelerations, and the third subject has idiosyncratic deep breaths followed by a ballooning pattern in the respiratory envelope in each run. To the extent that pseudomotion or real motion accompany breathing, motion estimates should share many of these properties.

**Figure 3:**
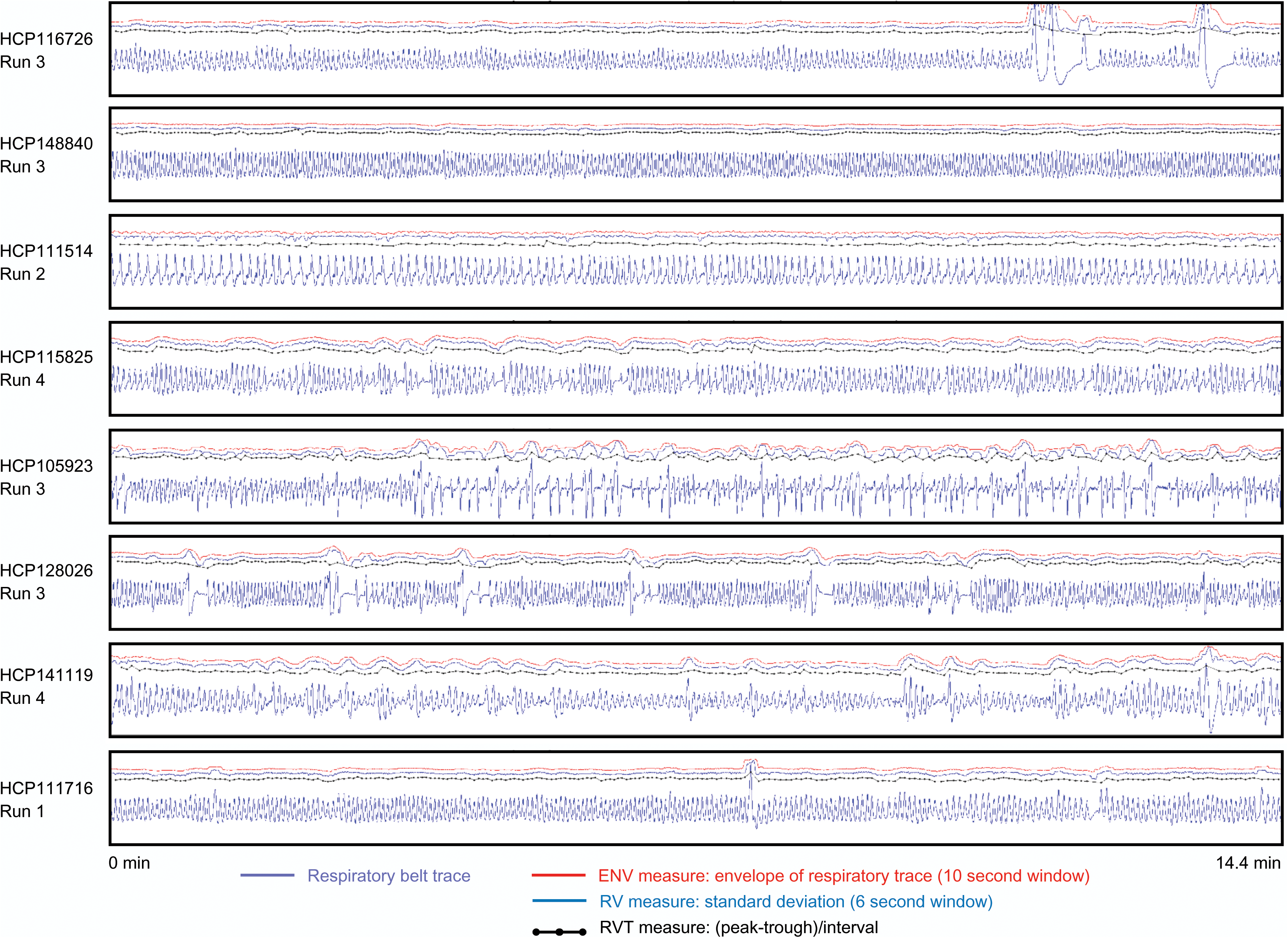
Examples of respiratory traces and derived measures. For 8 scans from 8 subjects of the HCP dataset, respiratory traces and derived ENV, RV, and RVT measures are shown. Each scan lasts 14.4 minutes. Note the wide variety of rate and depth of breathing within a scan and across subjects, the varied shapes of the raw respiratory belt waveforms, as well as the variety of pauses and very large breaths.

**Figure 4:**
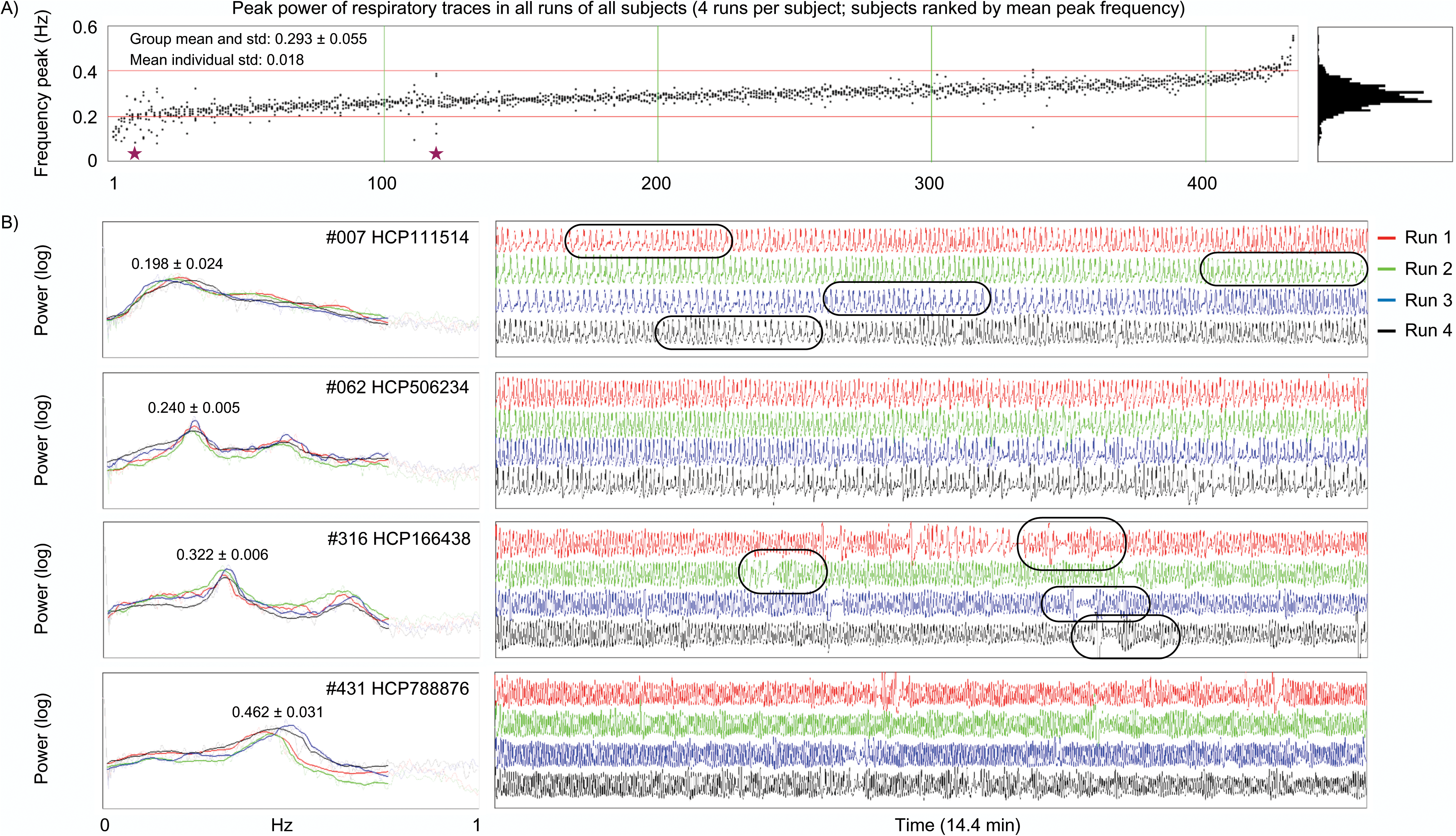
Frequency content of respiratory traces. A) The peak in power spectra of respiratory traces of each run of each subject, ordered by subject mean peak value. Most runs have peak power between 0.2-0.4 Hz. B) Example subject power spectra and respiratory traces. Spectra tend to be similar within a subject across runs, reflected in narrow variance in peak location in single subjects relative to variance across the group. See Figure S1 for examples of dissimilar spectra within individuals across runs (the subjects with purple stars in (A)). Note that many spectra exhibit rather narrow peaks (e.g., the last 4 examples of (B)) whereas in some subjects a rather broad range of respiratory rates are found within a run (e.g., the broad peaks of the first example of (B)).

The power spectra of respiratory and position traces contain informative similarities and differences (spectra of all subjects can be seen in online movies^2^). Spectra of all runs are shown for 8 individual HCP subjects in Figure 5, with clear representation of respiratory frequencies in the position traces. Additionally, there is a low-frequency peak around 0.10-0.15 Hz in the position traces that is not present in the respiratory traces (red ovals). When power spectra from all runs of all HCP subjects are shown as a heat map, ordered by peak respiratory frequency (Figure 5B), three properties are evident. First, the peak frequencies of respiration and its harmonics map onto the peak frequencies of position traces. Second, there is a position frequency peak around 0.12 Hz in virtually all subjects that is not present in respiratory traces, corresponding to the red ovals above. Third, there is a position frequency peak around 0.55 Hz in virtually all subjects that is not present in respiratory traces. The narrowness of the 0.55 Hz peak suggests that it is an artifact, and this will be confirmed below, but the 0.12 Hz peak has interesting properties that will be explored further. Figure 5C plots peak respiratory frequency against peak head position frequency, with excellent correspondence above ∼0.20 Hz. Below ∼0.2 Hz the true respiratory peak begins to merge with the 0.12 Hz peak and the correspondence weakens. The respiratory traces have not undergone any filtering that would influence signals at 0.10-0.15 Hz, but to ensure that the outlier replacement and 1-second blurring procedures used to prepare the respiratory traces did not somehow obscure a 0.12 Hz peak, identical analyses were performed on the raw respiratory belt traces that had undergone no processing, and these analyses yielded no evidence of a 0.12 Hz peak, or of a 0.55 Hz peak (data not shown). When analyzed similarly, pulse oximetry traces yielded no evidence of a 0.12 Hz peak (data not shown).

**Figure 5:**
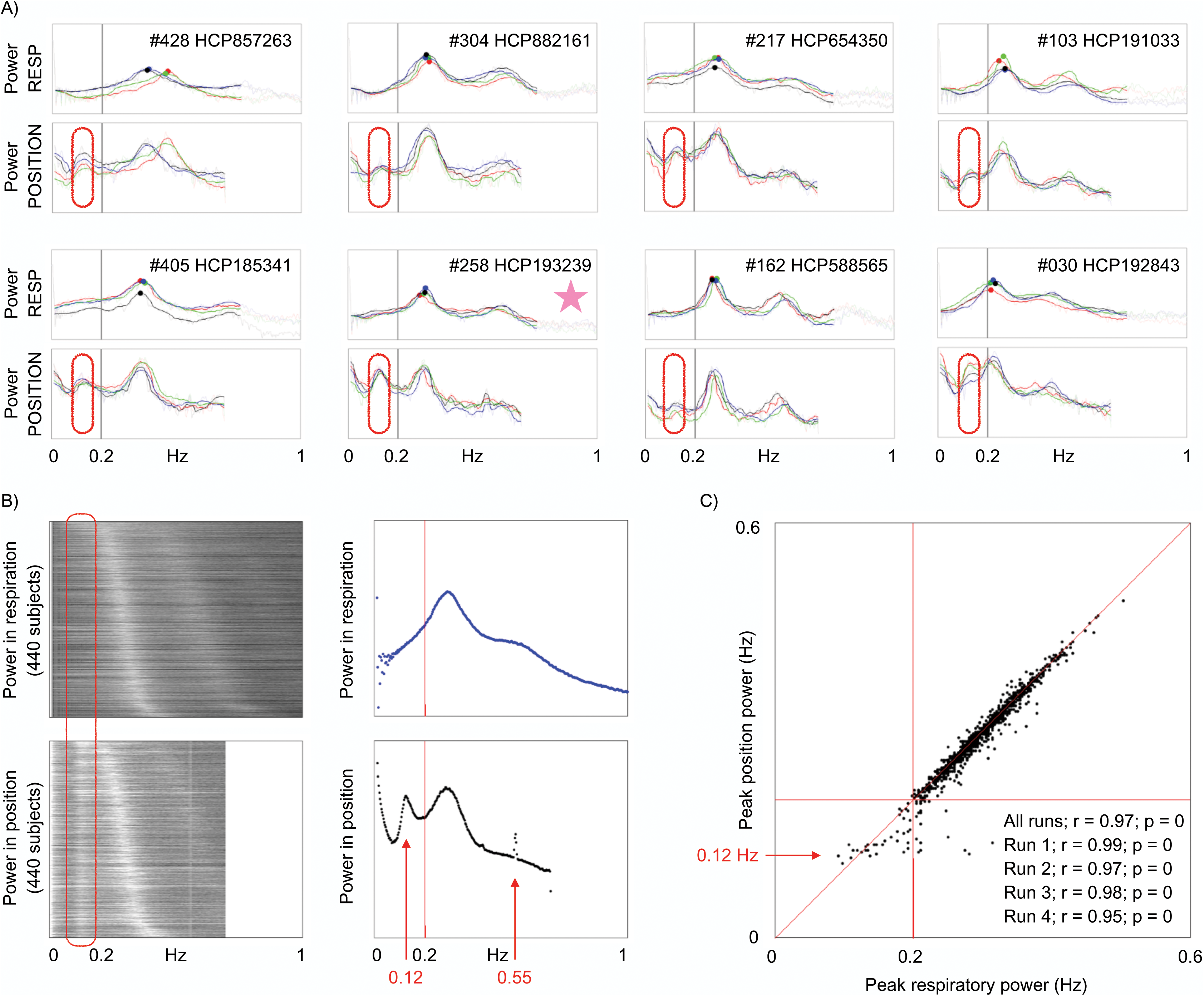
Comparison of respiratory frequency spectra with head position frequency spectra. A) Power spectra of 8 subjects are shown for respiratory and position traces (X realignment parameter); the 4 colors denote the 4 runs. Ovals circle a peak in position spectra not present in respiratory spectra. B) Heat map of spectra from all runs of all subjects, ordered by peak respiratory frequency. Scatter plots are means across subjects. C) Frequency of peak power in respiration plotted against frequency of peak power in head position. Correlations of peak frequencies across subjects are inset. The starred subject is examined in Figure 6.

### Identifying a low-frequency respiratory pseudomotion

Although the low-frequency 0.12 Hz peak in position traces is not in respiratory traces, it is a respiratory phenomenon. This fact is appreciable in single subjects. Figure 6 shows data from the starred subject of Figure 5A (this subject was chosen because it displays considerable amplitude at the low-frequency peak, see the 0.12 Hz peak height in Figure 5A). At top, respiratory traces of all runs are shown, with red ovals placed to encircle numerous deep breaths. Below, the X position traces are shown, with high-amplitude low-frequency waves emerging at the times of the deep breaths (see identically placed ovals). The third panel illustrates variance removed from position traces by a 0.08-0.18 Hz stopband filter, confirming that these waves are the ones that constitute the low-frequency peak. DVARS traces indicate that there are usually image abnormalities at these times. Thus, the low-frequency X position changes are associated with deep inspirations.

**Figure 6:**
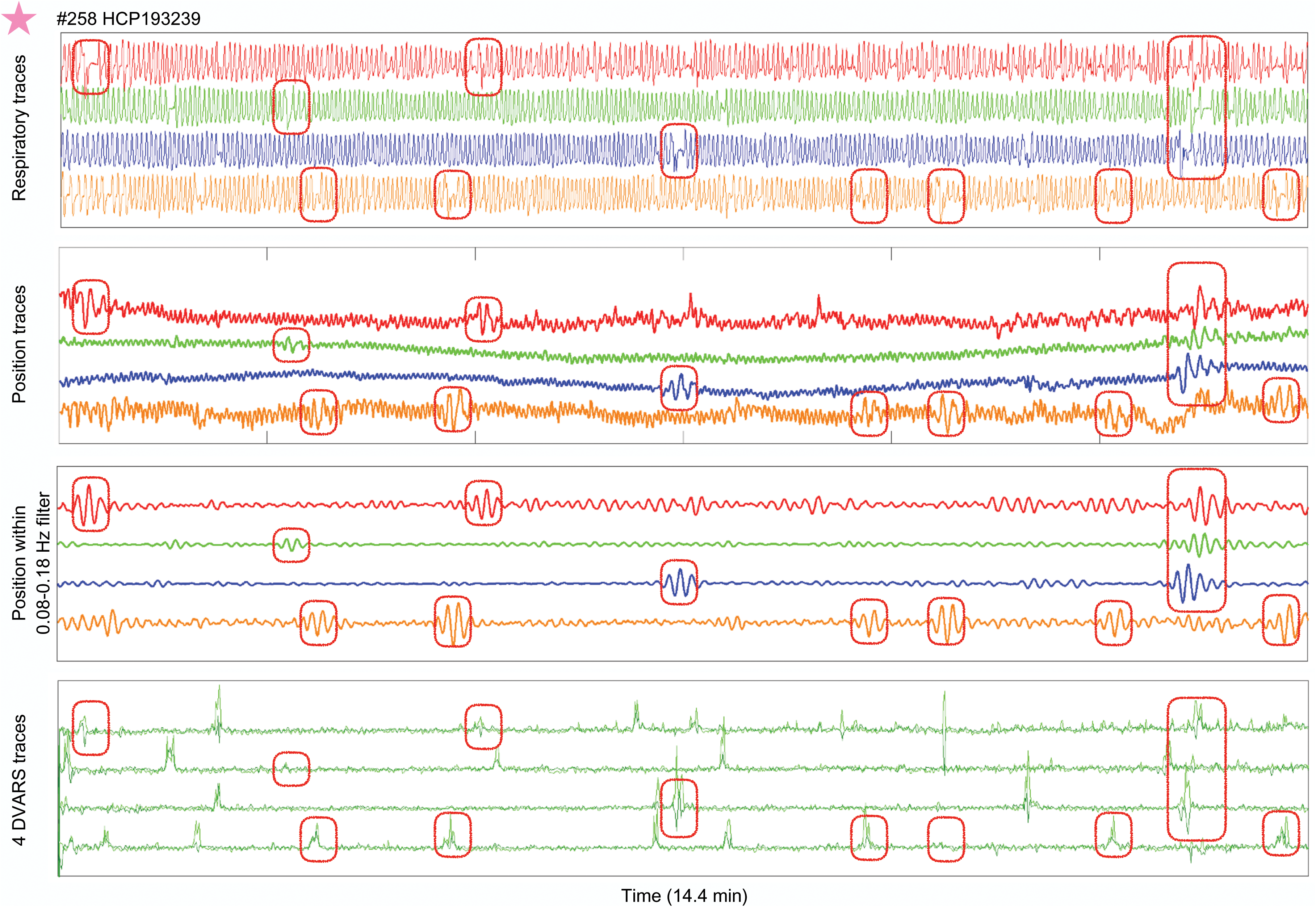
Low-frequency position waveforms emerge during deep breaths. For the starred subject of Figure 5, respiratory traces, position traces, and DVARS traces of all runs in their entirety are shown. Instances of deep breaths highlighted are in red ovals. Low frequency waves in position traces occur during these breaths, and a stopband filter spanning 0.08 Hz-0.18 Hz removes the variance in the third panel, confirming that these waves constitute the low-frequency peak of the data. This same subject is shown again in Figure 8.

Because subjects are scanned lying supine, the occiput is the natural pivot of the head due to movement of the chest, and respiratory motion should most prominently appear in the pitch rotation estimate and the superior-inferior translation estimate. However, the parameter used for Figures 5 and 6 is the left-right (X) parameter. Position parameters are coupled in rigid body realignment in ways fundamentally dependent on the origin of the coordinate system. Thus, one could posit that deep breaths cause some kind of true low-frequency motion, and the left-right parameter changes are due to coupled “bleeding” of motion from one parameter into another. There is, however, another explanation. Expansion of the lungs can shift the B0 field, which can manifest as apparent motion in the phase-encode direction despite the head remaining still, the phenomenon called pseudomotion (Brosch et al., 2002; Durand et al., 2001; Raj et al., 2001). The left-right direction is also the phase encode direction of the HCP data, and thus both true motion and pseudomotion could explain the low-frequency waves.

To help disambiguate these possibilities, several fast-TR datasets with different phase encode directions were obtained, and power spectra of all 6 position estimates of these datasets are shown in Figure 7. A low-frequency peak appears in each of these datasets, but only in the phase-encode direction (red arrows). The frequency of the low-frequency peak scales with a sequence parameter: the MyConnectome, Cornell, and HCP datasets have respective TRs of 1160, 766, and 720 ms, and peak frequencies of 0.08, 0.11, and 0.12 Hz. In other words, a TR 1.5 times faster yields a low-frequency peak approximately 1.5 times faster. Thus, these low-frequency waves in the phase encode direction are consistent with pseudomotion resulting from B0 shifts with deep breaths. The fact that this in-plane translational motion does not “bleed” into other parameters during realignment is informative because roll effects might have also been expected due to inferior brain tissue being closer to the lungs. By contrast, the peak respiratory frequencies of the HCP data appear in all position parameters, consistent with true head motion occurring in multiple parameters as the head pivots about the occiput and then “bleeding” into all position estimates via the rigid body assumptions. Whether this presumably true respiratory motion is accompanied by pseudomotion (in only the phase encode direction) will be examined below.

**Figure 7:**
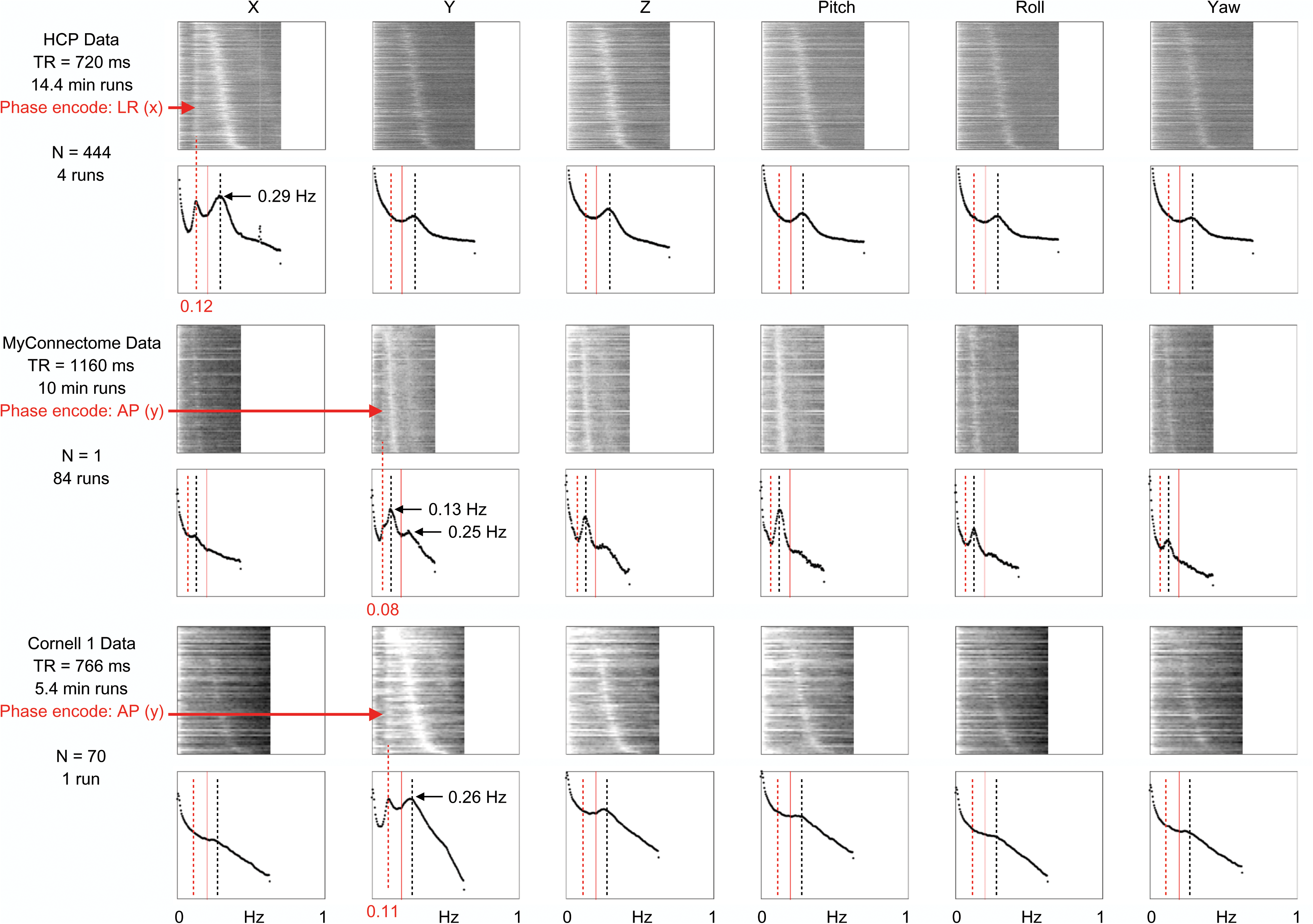
Low-frequency position waveforms occur only in the phase-encode direction. For 3 fast-TR datasets with 2 different phase encode directions, power spectra of position estimates are shown for all scans of all subjects. Each dataset contains a low-frequency peak (or shoulder) seen only in the phase-encode direction (red arrows), whereas frequencies typical of respiratory frequencies are seen congruently in all parameters. Note the respiratory harmonics of 0.13 Hz and 0.25 Hz in the MyConnectome dataset, visible because all 84 scans are of the same subject (whereas the other datasets sample dozens to hundreds of subjects and blur a variety of peak frequencies).

A fast-TR developmental dataset gathered on the same scanner as the Cornell 1 dataset yields a further interesting finding. In 11 children and teenagers scanned twice for 4.8 minutes with TR of 800 ms, typical respiratory frequencies are observed in position power spectra in all directions, but there is no low-frequency peak in the phase encode direction (Figure S2). The apparent absence of the low-frequency pseudomotion in the Cornell 2 dataset may be due to the relatively small lungs and bodies of these children, which should produce less perturbation of B0 fields during breaths than adult bodies produce, or, if the low-frequency pseudomotion relates to a physiological phenomenon, perhaps this process emerges over development. These findings suggest that different kinds of motion may manifest over development for technical (or biological) reasons in addition to the tendency of children to be restless and active in the scanner.

A final comment on these findings is that only in the HCP data is a 0.55 Hz peak in position spectra seen, making the scanner, sequence, or environmental artifacts specific to this dataset the most likely explanation of this peak. Conceivably, the low-frequency pseudomotion could be related to a low-frequency physiological phenomenon, such as the Mayer waves that occur at similar frequencies (Strik et al., 2002), but we see no obvious mechanism by which such arterial pressure waves would produce effects only in the phase encode direction.

### Using unprocessed images to disambiguate pseudomotion from real motion

Pseudomotion can be distinguished most clearly from real motion in one of two ways: by demonstrating a pseudomotion in only the phase encode direction in unprocessed images, and by demonstrating a real motion in a non-phase-encode direction in unprocessed images. To describe motion in unregistered, unprocessed, scanner space images, one can calculate the centers of mass of the entire image, yielding motions in the X, Y, and Z directions (rotations may or may not influence the center of mass). A number of unprocessed HCP and MyConnectome datasets were obtained, and center of mass traces were calculated and inspected. Readers may view data from all scans online^3^, and data from several scans are shown in Figure 8. In HCP scans there are visible respiratory fluctuations in the phase encode (left-right), frequency encode (anterior-posterior), and slice stack (superior-inferior) directions. MyConnectome scans also exhibited respiratory-appearing motion in all 3 directions (respiratory records were not acquired with MyConnectome datasets). These results indicate that much of the constant respiratory motion is real motion because it is not limited to the phase encode direction.

**Figure 8:**
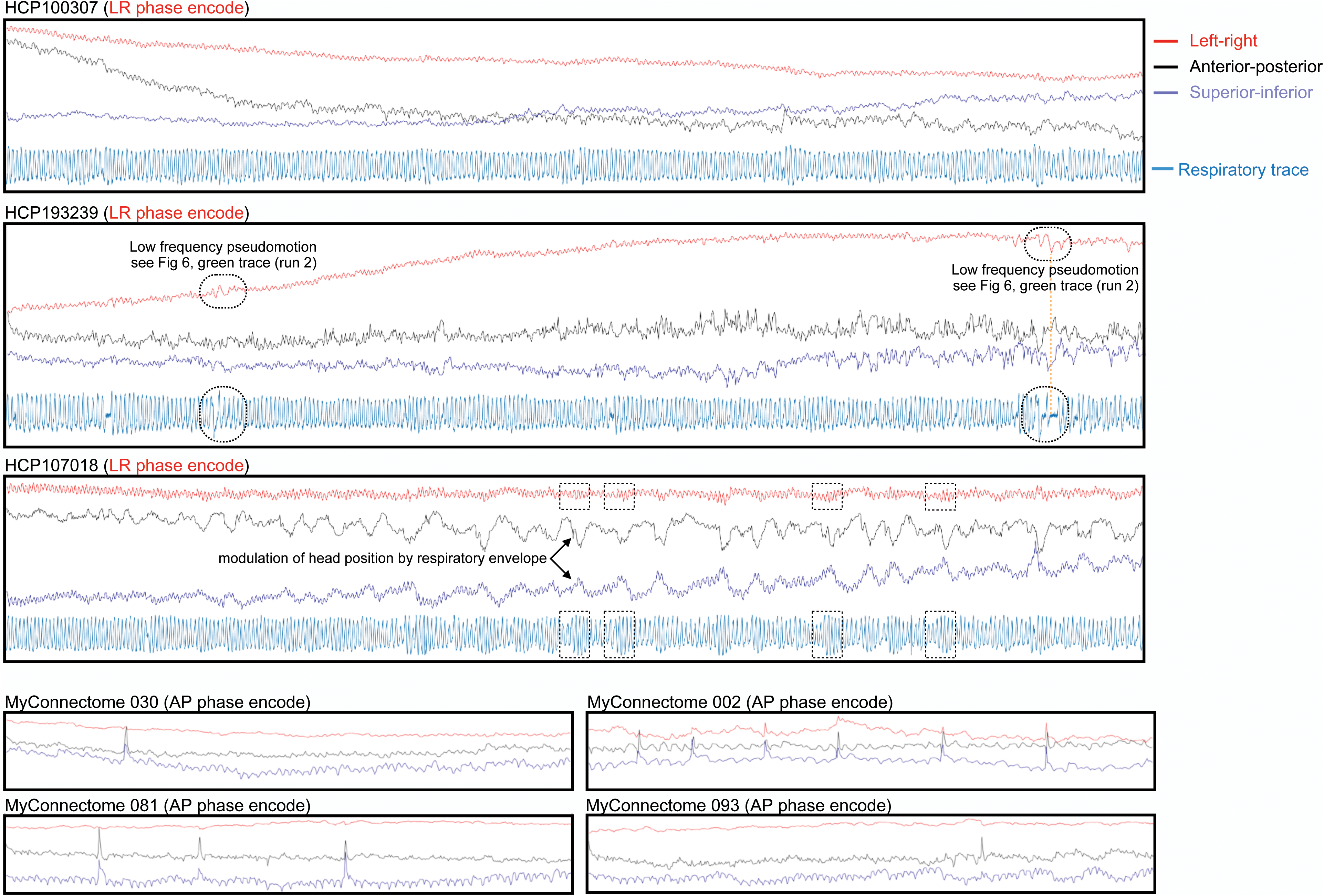
Plots of the center of mass of unprocessed scans from the MyConnectome and HCP datasets. Respiratory fluctuations are evident in all 3 directions in scanner space, but especially in the superior-inferior direction (slice stack) and the phase encode directions.

The head will be variably placed within scanner space, but, assuming a reasonably aligned placement, one would expect respiratory nodding to substantially appear in the superior-inferior direction, to partially appear in the anterior-posterior direction, and to rarely appear in the left-right direction. Compatible observations are made in the unprocessed images. First, constant respiratory fluctuation in the superior-inferior direction is seen in all scans. Second, constant respiratory fluctuation in one or both of the left-right and anterior-posterior directions is seen in most scans, and is accentuated in the phase encode direction. Re-examination of Figure 7 yields congruent information: Z and pitch parameters have the strongest respiratory power in all datasets, as does whichever parameter is the phase encode direction. This augmentation of respiratory power in the phase encode direction presumably reflects the contribution of pseudomotion at the respiratory rate. These patterns accord with the findings in (Raj et al., 2001), in which left-right oscillations were absent in acquisitions with anterior-posterior phase encode direction but were prominent in left-right phase encode acquisitions, whereas anterior-posterior oscillations were present regardless of phase encode direction but were stronger with anterior-posterior phase encode acquisitions. Similarly, the mannequin constructed by Brosch and colleagues to investigate respiratory pseudomotion yielded only phase encode effects (Brosch et al., 2002).

Pseudomotion and real motion can be dissociated from one another in time. Under normal circumstances, when air is moving into and out of the lungs, both real head motion and pseudomotion could scale with deeper breathing. An example of pseudomotion scaling with breathing depth is shown in the boxed portions of Figure 8 (real vertical head motion in these instances is present (e.g., in the vertical orientation) but is not visibly changing). However, during apneas, dissociations can occur. If an apnea is central (i.e., there is no respiratory drive and no ventilatory effort), both pseudomotion and respiratory motion should essentially cease. But if an apnea is obstructive (i.e., soft tissues impede air flow), pseudomotion should cease but respiratory efforts may continue and even increase. Instances of both of thes ephenomena can be found in the 10 subjects we examined, and examples are shown in Figure 9. Note that the inspiratory efforts can cause large vertical and anterior-posterior motions during partial or complete obstruction as the subject attempts to breathe against resistance and/or restore air flow.

**Figure 9:**
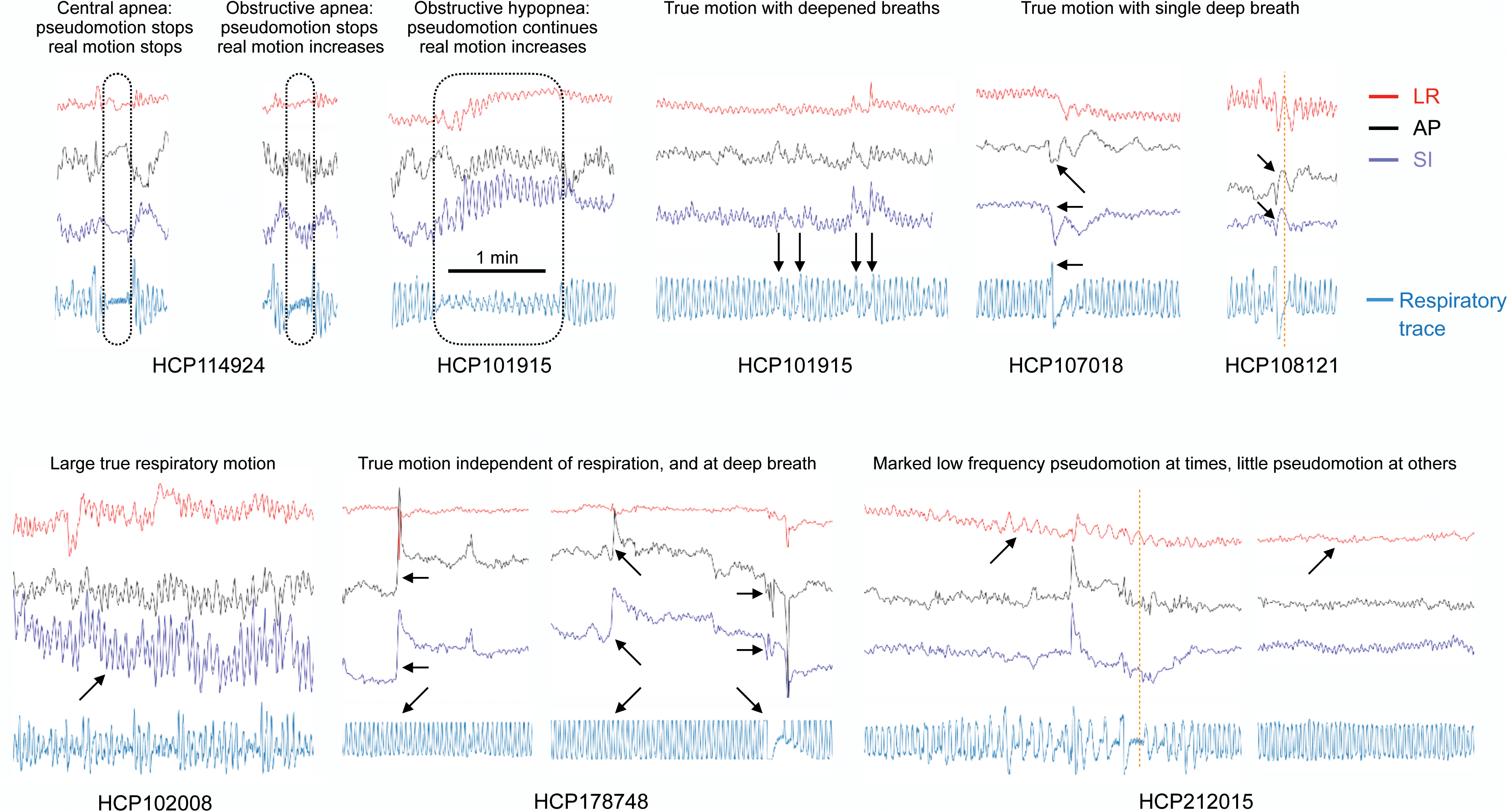
Examples of motion and pseudomotion. Small excerpts from several scans are shown, illustrating central apnea, obstructive apnea, obstructive hypopnea, true step motions occurring with substantial changes in breathing depth, true motions occurring with deep breaths, marked real respiratory motion, real motions without any respiratory contribution, low-frequency pseudomotion in the absence of respiratory motion (orange dotted lines), and variable appearance of pseudomotion within a subject. LR: left-right; AP: anterior-posterior; SI: superior-inferior.

The occurrence and magnitude of true respiratory motion is likely to depend on characteristics of the subject (and on the steps taken to physically restrain the head). For example, subjects with asthma will exert themselves more to draw breaths, and if partial airway obstruction occurs as a subject relaxes or sleeps, snoring or inspiratory efforts will ensue at the breathing frequency. That the amplitude of constant real respiratory motion can become very large is both self-evident and demonstrated in Figure 9.

The occurrence and magnitude of pseudomotion will largely depend on scan parameters (e.g., phase encode direction, field strength), and the size of lungs and the body. Additionally, the magnitude of pseudomotion can fluctuate, as demonstrated in the examples above of apnea and deepened breathing, but these fluctuations appeared limited to perhaps two or three-fold the eupneic fluctuations. We saw no instances of left-right pseudomotion amplitude increases comparable to the large increases in real vertical fluctuations shown in Figure 9; when very large oscillations occurred, they only appeared in anterior-posterior and vertical orientations. The largest amplitude changes in pseudomotion are seen during very deep breaths, which, by their nature, occur more slowly than typical respirations. Deep breaths also produce the distinct low-frequency phase encode phenomenon described above. That such low-frequency fluctuations are not the same as ongoing respiratory pseudomotion can be seen in several instances in Figures 8 and 9 where orange dotted lines mark reversals in phase-encode position occurring at times of apnea or unchanging respiratory direction.

Beyond the constant respiratory motion and pseudomotion, large changes in vertical and anterior-posterior positions that followed the respiratory envelope (but not the respiratory rate) were often seen (see black and blue traces at the boxed times of Figure 8). The magnitude of these changes is generally considerably larger than that of the constant respiratory oscillations of the phase encode direction. Other examples of real step motions in all three directions at deep inspirations are seen in Figure 9, as are examples of real motions in all three directions that appear completely unrelated to respiration.

Collectively, these observations indicate that the head is in constant motion due to respiratory cycles, that pseudomotion augments this constant motion in the phase encode direction, but that much constant motion and most major motions are not attributable to pseudomotion.

### Re-presenting motion in fast-TR data

In a contemporary fast-TR dataset like the HCP data, based on the above observations, investigators may encounter at least seven kinds of motion, five of them related to respiration: 1) constant real respiratory motion in the form of head nodding most evident in vertical position and pitch, which can be very large at times; 2) constant pseudomotion at the same respiratory rate as real motion, occurring only in the phase encode direction; 3) punctate real motions occurring at times of very deep breaths; 4) a low-frequency pseudomotion in only the phase encode direction following very deep breaths; 5) slow modulation of vertical and anterior-posterior head position by the respiratory envelope; 6) punctate real motions unrelated to respiration; and 7) periodic or transient artifacts such as the 0.55 Hz motion of the HCP data. The existence of distinct kinds of motion raises interrelated questions: how well can the kinds of motion be separated, are effect sizes meaningful if different kinds of motion are separated, and what purposes could be served by separating kinds of motion?

In the case of pseudomotion, it seems self-evident that one might wish to separate “apparent” from “true” motion for a variety of purposes (e.g., one is within-plane motion and largely a respiratory epiphenomenon, the other is largely through-plane motion with different physical consequences for fMRI signals).

The low-frequency pseudomotion can often be isolated by frequency filters, since most subjects have respiratory frequencies well above the low-frequency pseudomotion. Figure 10 illustrates filtering the phase-encode direction position parameter to suppress low-frequency pseudomotion in two representative subjects (compare gray and black X position traces, and the corresponding gray and black FD traces). Slight changes in both traces are seen after filtering. The FD effects are small because low-frequency pseudomotion is typically prominent during only a small fraction of the scan and affects only one of six position parameters. Additionally, though low-frequency pseudomotion effects are large amplitude in some subjects, this is not true of most subjects (see the relative peak heights in spectra of Figure 5). Across subjects, filtering out low-frequency pseudomotion results in mean FD correlated at r = 0.999 with original FD, with magnitude of 98.9% of original magnitudes (Figure S3). Practically, for purposes of motion estimation, filtering out such pseudomotion has little effect in most subjects. The largest effects occur in subjects with low respiratory frequencies, which is predictable because in such subjects the low-frequency filter (here, a stop band of 0.08 – 0.18 Hz) removes some of the true cyclic respiratory motion. Although removing low-frequency pseudomotion has little impact on motion estimates in most subjects, the isolation of low-frequency pseudomotion via filtering could potentially serve as an index of deep respiration in data lacking respiratory records.

**Figure 10:**
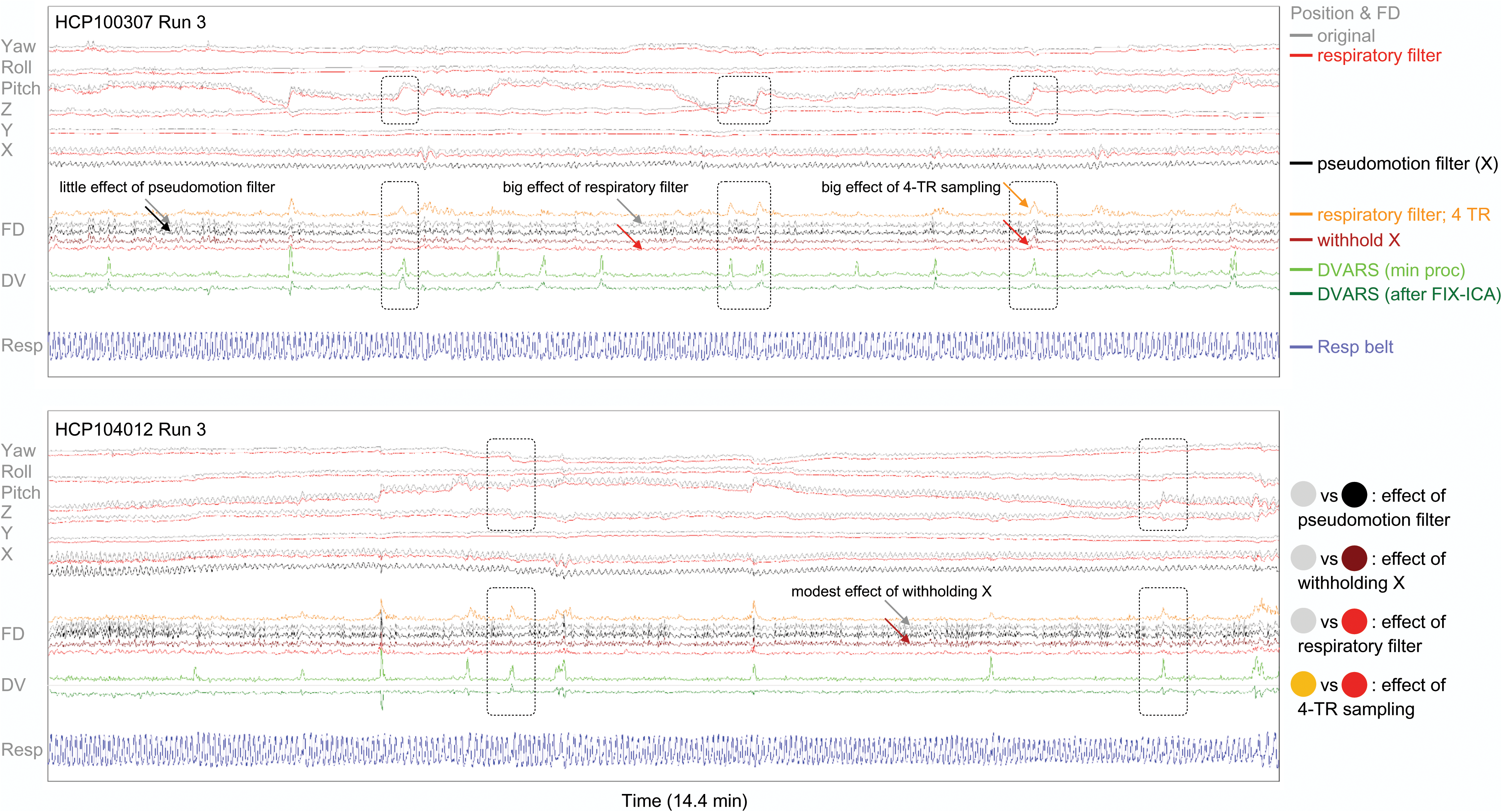
Modifications of motion estimates. Runs for two subjects are shown. At top, the 6 position estimates: gray traces show the original estimates, red traces show estimates after removing 0.2-0.5 Hz respiratory frequencies, and the black X trace represents the phase encode direction after removing 0.08-0.18 Hz pseudomotion frequencies. In the middle, FD derived from the three kinds of position traces at top are shown. Filtering out pseudomotion has little effect on FD (compare gray and black FD traces) whereas filtering respiratory frequencies has a large effect (compare gray and red traces). Withholding the X parameter has a modest effect. At bottom are the respiratory trace and DVARS traces for reference. Note that several motions are apparent in the raw position estimates, with corresponding DVARS abnormalities (boxes). However these motions are lost amidst the respiratory noise in the gray and black and maroon FD traces, and are also not prominent in the red traces. When a 4-TR differential in motion is calculated (orange FD trace), true head motions stand out from the rest of the trace and match what is seen in DVARS.

A different way to eliminate pseudomotion would be to omit the phase encode direction entirely from motion calculations. This approach is simple, but has a major practical constraint: though one might be relatively comfortable omitting a left-right phase encode position parameter from motion calculations in HCP data (since little true motion is expected in this direction), one would probably hesitate to eliminate an anterior-posterior phase encode parameter in a different dataset (since much true motion occurs in this orientation). For illustrative purposes, FD calculated without the left-right parameter is shown for the subjects in Figure 10 (compare gray and maroon FD). Because there is constant real motion at the respiratory rate in multiple parameters of comparable (or greater) magnitude than the pseudomotion in the phase encode parameter, ongoing respiratory motion remains a prominent effect in the trace. Across subjects, omitting the phase encode direction results in mean FD correlated at r = 0.921 with original FD, with magnitude of 66.7% of original magnitudes (Figure S3). Because this approach seems unlikely to be used in datasets with anterior-posterior phase encode orientations (the bulk of our own data, and much other data), we do not pursue this avenue further.

The dominant respiratory frequency is present at all times in the scan and affects all position parameters. Filtering out those frequencies has a large effect on head motion (compare gray and red traces in Figure 10). Filtered position estimates are “smoothed” out with relative preservation of step changes. The filter in Figure 10 has a 0.2 – 0.5 Hz stop band, and across subjects this filter yields mean FD with magnitude 51.9% of the original magnitudes, correlated at r = 0.819 with original FD (Figure S3). Some comments should be made about filter selection and application, but prior to discussing these issues, it is helpful to consider what purpose is served by removing respiratory motion.

There are two main reasons to separate constant respiratory motion from episodic or gradual motion. The first reason is conceptual: different neural phenomena and behavioral or biological predispositions may underlie constant respiratory versus episodic/gradual motion. For example, constant respiratory motion may relate to body habitus, whereas episodic/gradual motion may more closely reflect restlessness, impatience, fatigue, or anxiety. Separating kinds of motion can help identify such distinct relationships. The second reason to filter out respiratory motion is practical: motion traces become more visually useful and more congruent with other data quality traces once constant respiratory motion is suppressed.

In fMRI datasets with TRs of 2-4 seconds, it is customary to see FD traces with well-defined, flat “floor” values from which brief, clear excursions are seen during motion, and these FD traces closely resemble DVARS traces and other data quality traces (Power et al., 2012; Power et al., 2014; Satterthwaite et al., 2013; Yan et al., 2013). Multiple authors have noted that the HCP FD traces do not have these properties, and, instead, there appears to be constant motion in the traces often without clear excursions even when motion is known to occur (Burgess et al., 2016; Glasser et al., 2018; Power, 2017). There are two reasons for such properties. First, constant, cyclic, respiratory motion used to be aliased with slower fMRI sampling rates but is not aliased in HCP data due to the faster sampling, and the head is truly in constant motion in the images. Second, and separately, episodic or gradual shifts of head position that would appear large if measured every 2-4 seconds become subdivided in fast TR data to such an extent that the motion can “disappear” into the cyclic respiratory motion. Two steps can be taken to counter these two phenomena. First, respiratory filtering can suppress much of the cyclic motion. Second, calculating FD over multiple TRs rather than a single TR can “undo” the subdivision of episodic or gradual motion. To illustrate these steps, FD was calculated over intervals of 4 TRs (4 * 720 ms = 2880 ms), using the position estimates with 0.2-0.5 Hz frequencies removed, to yield the orange FD traces in Figure 10. These orange FD traces show large excursions from a floor value, at times when motion is visually evident in the position traces, during which times image abnormalities are indexed by DVARS spikes. These orange FD traces are visually useful in ways comparable to FD traces in traditional slow-TR datasets; they plainly reflect the changes visible in position estimates.

Returning to filter selection and application, there is no single “correct” way to treat or filter motion estimates to suppress respiratory motion, for the simple reason that respiratory rates vary widely across and within subjects. Some subjects have low respiratory rates that overlap with pseudomotion frequencies. Many more subjects have variable respiratory rates that sometimes drop to low frequencies for portions of scans. An example is in Figure 10 at the end of the bottom scan, where the respiratory rate drops dramatically in the final 30 seconds of the scan, and respiratory motion starts to escape the respiratory filter (red position traces), causing a continuous elevation of FD even after filtering (see red and orange FD traces). When a subject’s frequency peak is narrow (i.e., the subject exhibits a relatively constant periodicity of breathing), a rather narrow stop band filter could suppress much respiratory motion. But if the frequency peak is broad (i.e., there are times of slow and of fast breathing, e.g., the bottom panel of Figure 10), then a wider stop band is needed to suppress respiratory motion.

Faced with these characteristics, an investigator can apply a uniform filter intended to suppress much but not all of the respiratory motion (the choice made here), or apply a filter tailored to each subject’s respiratory characteristics. The width of the stop band interacts with the ability to localize motion in time. Step changes and brief spikes in motion have broadband frequency characteristics, and, as stop bands widen and remove more frequencies, the ability to represent rapid position changes diminishes. This fact is seen in two forms in Figure 10: the lower slopes of red filtered position traces compared to gray unfiltered position traces at step changes, and the relatively small sizes of peaks in the red filtered FD traces. The boxed portions of Figure 10 show several instances where motion is evident in the original position traces, but is difficult to discern in the red filtered FD traces until drawn out by calculating 4-TR differentials shown in the orange FD trace. An additional factor in filter selection is that the broader the stop band, the fewer the degrees of freedom in the residuals. We chose our filter to be uniform across subjects in order to avoid adding further complexity to the paper. The filter parameters were chosen to suppress most respiratory motion in most subjects, knowing that respiratory motion may persist if breaths are sufficiently slow, but balancing this consideration against wanting clear peaks of motion. Interpretations of our motion-related findings below will bear these caveats in mind (and, at times, we only examine subjects with peak frequencies within the respiratory stop band (i.e., above 0.2 Hz) to minimize confounds of unsuppressed respiratory motion).

Finally, we should note that if an investigator possesses respiratory records, those records (or treatments of the records) can be used as nuisance regressors in the motion traces. However, the relationships that are so visually evident between respiratory records and motion traces can be complicated to capture statistically in a regressor: respiratory records often hit ceiling and floor values during deep inspirations and exhalations, and position does not tend to scale linearly with belt values during deep breaths; obstructive apneas dissociate motion from respiratory fluctuations; and position estimates contain a variety of step changes and gradual shifts not present in the respiratory traces. For these reasons, regressors from belt traces can show markedly different success at removing respiratory motion across scans and even across single position parameters in a scan. Figure S4 shows instances of success and failure. With regard to identifying or removing respiratory motion, respiratory records are probably mainly useful for independently establishing respiratory frequencies (which, for example, could be used to guide a wavelet-based removal of respiratory motion).

The net effect of filtering out respiratory frequencies and calculating motion over multiple TRs is shown in Figure 11 (similar plots are shown online for all subjects^4^). The original motion estimate shows nothing informative beyond seemingly constant motion. Once respiratory frequencies are suppressed and a 4-TR differential is calculated, motion estimates show modest but distinct episodes of motion corresponding to the visible changes in position estimates, and to changes in the centers of mass in anterior-posterior and superior-inferior directions. DVARS spikes are present at these times, but DVARS dips are largely absent. These images demonstrate how motion traces can be modified to become informative about events in a scan.

**Figure 11:**
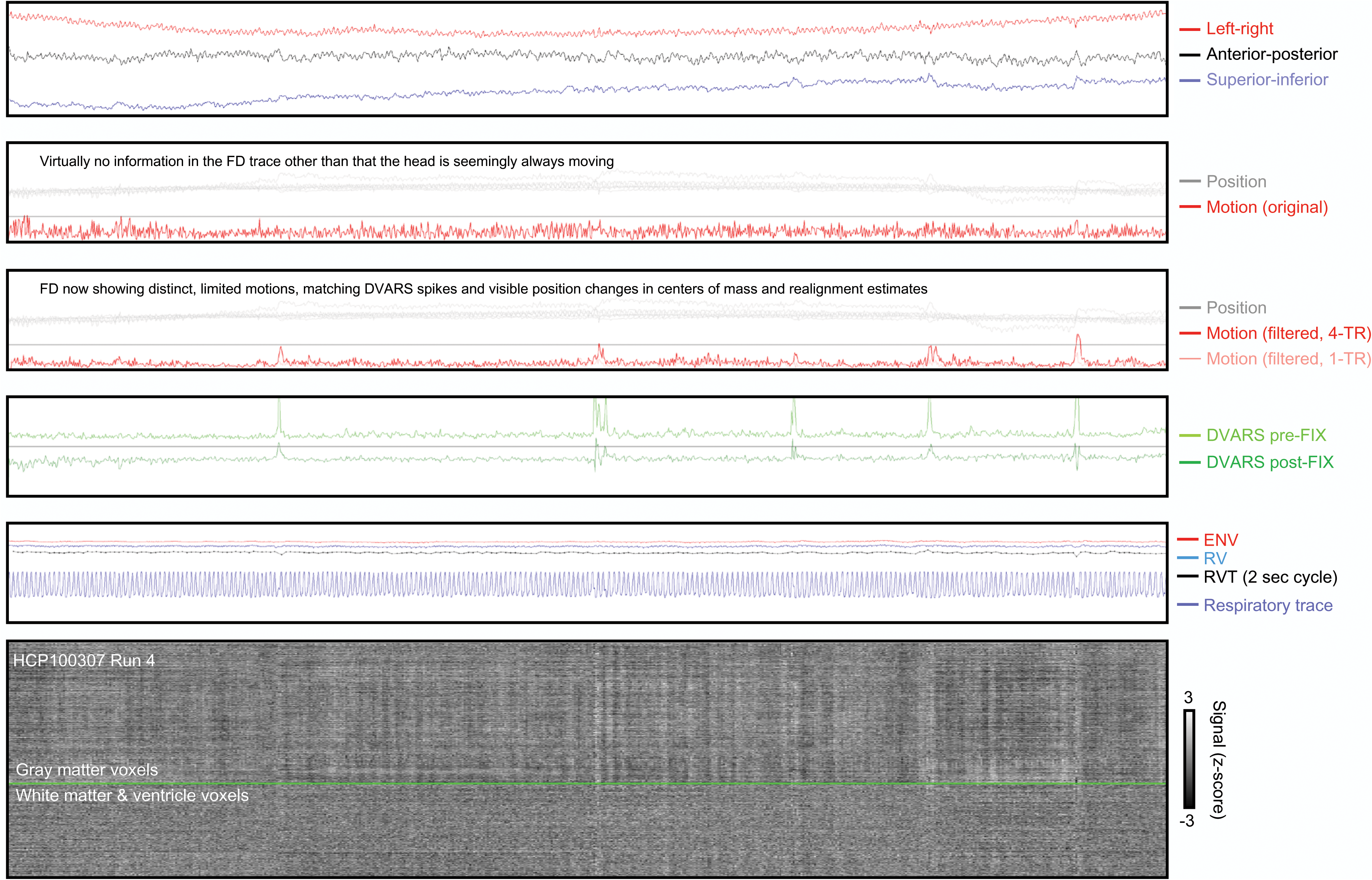
Illustration of the utility of filtering motion estimates and sampling motion over 2.88 seconds instead of 720 ms. Panels are as in prior figures: at top, centers of mass of the unprocessed image, next the original position and motion estimates, next the filtered motion estimate using 4-TR sampling is shown in red, with a faint trace showing the 1- TR version. Bottom panel shows signals of the minimally processed image.

### Links of motion to body habitus and respiration

In the center of mass data, we encountered examples of deep breaths causing spikes of motion, and also more gradual shifts in position that corresponded to changes in the envelope of breathing. Measures that capture such respiratory events, like the envelope of the respiratory waveform (ENV, (Power et al., 2018)) or windowed variance in the belt trace (RV, (Chang and Glover, 2009)), should thus be associated with motion (note that depending upon the shape of the deep inspiration, the RVT measure of (Birn et al., 2006) may or may not mark the event, because this measure depends on timings of peaks and troughs in the trace, see Figure 3). In slow-TR datasets, variance in respiratory measures like ENV and RVT scales with mean motion (Power et al., 2018; Power et al., 2017b). We therefore sought to establish whether this relationship was also found in fast-TR data.

A strong association of summary motion and variance in respiration measures is found in the HCP data. The standard deviation of the ENV trace and mean FD correlate at r = 0.51 across subjects (p = 0; see scatter plot). This statistic is obtained using the mean of the summary values across each run (std ENV and mean FD_filtered,4-TR_), and using only a single subject per family, but it does not matter whether all subjects are used, whether only single runs are examined, whether also only subjects with respiratory frequencies over 0.2 Hz are examined, or whether also high-motion subjects are excluded from the calculation (Figure 12). If within-subject linear fits are calculated across the 4 runs between the standard deviation of ENV and mean FD, the betas are 0.40 ± 0.93 (p = 0 by one-sample t-test). A similar set of findings emerges from calculations using RV and RVT, both with the correlations (Figure 12) and with the across-subject betas: RV betas are 0.39 ± 0.74 (p = 0) and RVT betas are 0.41 ± 1.33 (p = 0). All of these tests converge on the finding that increased variance in respiratory measures scales with increased motion, both across subjects and within subjects. If FD with respiratory filtering only is used (FD_filtered_), the relationships above are weakened to approximately r = 0.35, and remain significant. If FD without respiratory filtering is used (FD_original_), permitting constant respiratory motions to overshadow other motions, the relationships are largely lost (Figure 12). These findings demonstrate a strong link between respiration and motion that exists beyond an effect of pseudomotion: the effect is strongest when constant respiratory cycle motion (including pseudomotion) is maximally suppressed, and is largely abolished when respiratory cycles are permitted to influence motion estimates.

**Figure 12:**
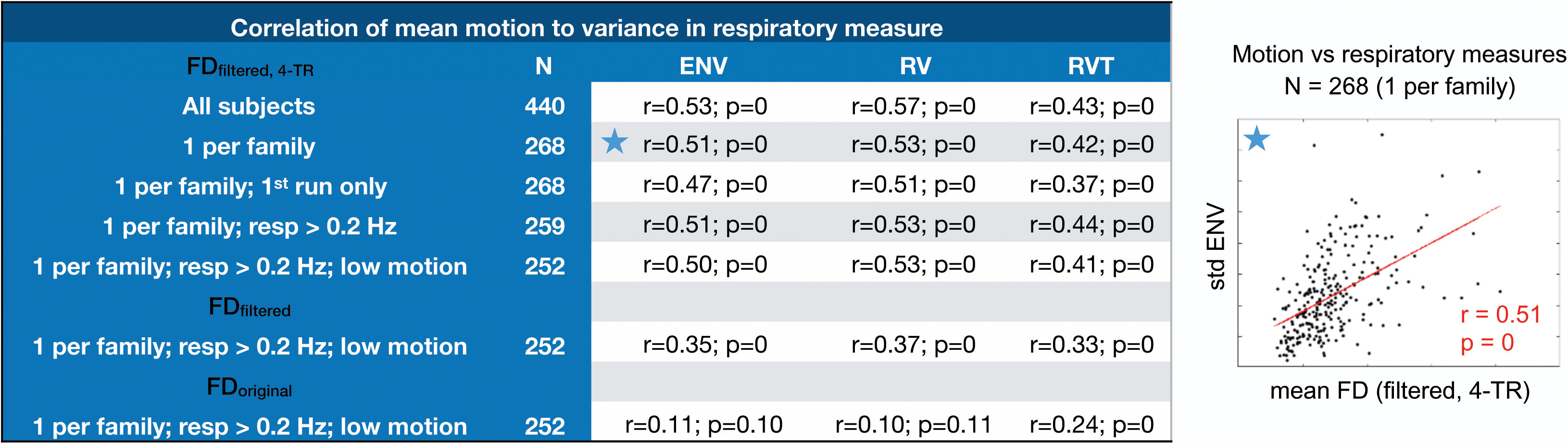
Relationship of summary respiratory measures to summary motion measures. Correlations between mean FD and standard deviations of ENV, RV, and RVT are shown, for several versions of cohort formation, and for several versions of FD. The scatter plot demonstrate the basis of the starred statistic of the chart.

Pseudomotion is suspected to arise as a function of changes in lung volume, due to both expansion of the lung cavity and displacement of the organs and soft tissues, but the relative contribution of each factor has been debated. In general, as BMI increases, tidal volumes and lung capacities decrease, whereas the work of breathing, minute ventilation, and respiratory rates all rise (Littleton, 2012; Luce, 1980). If expansion of the lung cavity is the important factor in causing pseudomotion, one might expect less pseudomotion in heavier subjects, and one might predict more pseudomotion in taller subjects, since lung volume scales strongly with height (Hepper et al., 1960). Either prediction could interact with respiratory rate, since shorter subjects breathe faster, as do heavier subjects (Chlif et al., 2009; Sampson and Grassino, 1983). In the HCP data, however, FD_original_ has no correlation with height, but correlates at r = 0.5-0.7 with weight or BMI (Figure 13, using 1 subject per family). When individual realignment parameters are examined, the amplitude of fluctuations in the left-right phase encode direction scales at r = 0.8 with BMI but not at all with height. Evidently, the soft tissues are the decisive factor in causing pseudomotion. Note that this special scaling of BMI with motion in the left-right parameter is not necessarily a reflection of how much pseudomotion contributes to overall BMI-associated motion; most precisely, it reflects how little left-right motion there is aside from pseudomotion.

**Figure 13:**
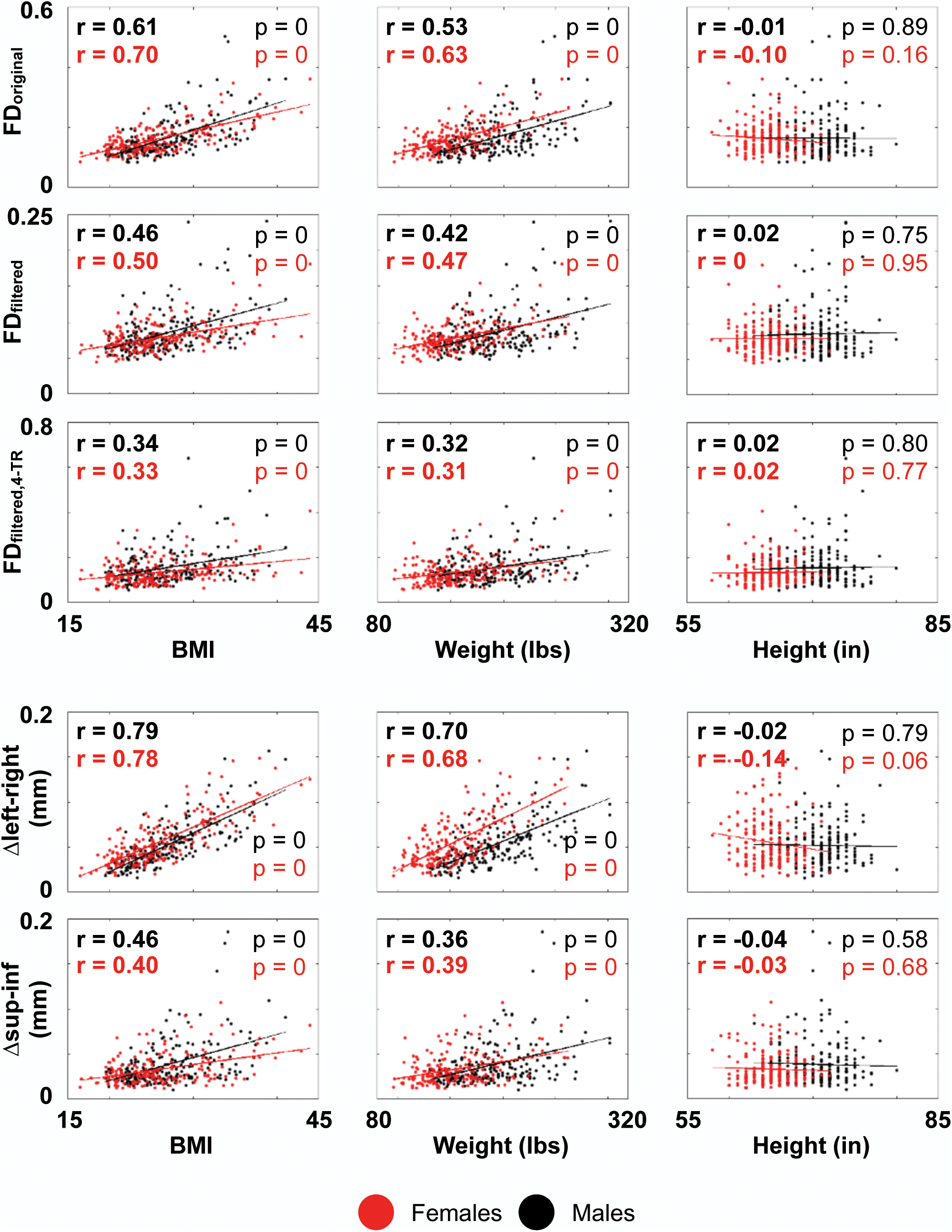
Relationship of summary motion measures to summary body habitus measures.

The scaling of BMI and motion could be due to the role of soft tissue mass in generating pseudomotion, but it might also be explained by increased minute ventilation and respiratory rates in heavier subjects. Figure S5 summarizes relevant findings. As expected, there is a modest correlation of BMI and weight to respiratory rate (BMI: r = 0.17 and 0.26 for males and females; weight r = 0.15 and 0.17). Respiratory rate correlated with mean FD_original_ in males at r = 0.14 and at r = 0.32 in females (likely due to the influence of the shorter subjects: in males height did not correlate with respiratory rate (r = 0.01) but in females, the shortest subjects around 5 feet tall had notably higher respiratory rates, causing a modest correlation of r = −0.22 between height and respiratory rate). The relationship of respiratory rate to motion was abolished in males and females after respiratory filtering of motion estimates. Thus, the explanatory power of respiratory rate for motion is much less than that of BMI, and the influence of respiratory rate disappears after filtering whereas BMI displays a persistent if weakened relationship to motion.

Given the persistent relationship of BMI to motion beyond influences of respiratory rate, we wondered if the respiratory variables that indexed motion in Figure 12 might mediate the persistent influence of BMI on motion, but there was no evidence for this (e.g., correlations of variance in ENV with BMI were r = 0 for males and r = 0.06 for females, and comparably low for weight and height). Different patterns of obesity exist, but generally, as BMI increases, fat deposits between the ribs and organs, reducing the flexibility of the trunk (Littleton, 2012). We speculate that this lack of flexibility may cause motions of all kinds to become more likely to be transferred to the neck and head.

### Heritability of respiratory and motion parameters

A prior study demonstrated in multiple datasets (including HCP data) that BMI and head motion (corresponding to FD_original_) are heritable traits, as is peak respiratory rate (Hodgson et al., 2017). In the HCP subjects studied here, 31 monozygous (MZ) and 23 (DZ) dizygous twin pairs are present (more subjects are twins, but without an accompanying twin under analysis). These numbers are not high, but permit preliminary descriptions of the heritability of respiratory and motion parameters studied in this paper. As mentioned earlier, respiratory power spectra are much more similar within-subject (r = 0.79 ± 0.15) than across all subjects (r = 0.44 ± 0.30). These observations extend to the twins: MZ pairs exhibit mean correlations between spectra of r = 0.64 ± 26, and DZ pairs exhibit mean correlations of r = 0.46 ± 0.34. As mentioned earlier, peak respiratory frequency is usually stable within subjects (standard deviation 0.018 Hz across 4 runs compared to 0.055 Hz across the group). As in (Hodgson et al., 2017), we find heritability of respiratory rate (MZ intraclass correlation coefficient = 0.48 (p = 0.002) and DZ ICC = 0.25 (n.s.), estimated H^2^ = 0.46, compared to H^2^ = 0.62 in (Hodgson et al., 2017)). Summary respiratory parameters such as variance in ENV are more similar within individuals than across the group (standard deviation within subject 0.048 vs 0.082 for the group), and are heritable (MZ ICC = 0.50 (p = 0.002) and DZ ICC = −0.14 (n.s.)). For variance in RVT, similar findings emerge: within-subject standard deviation of 0.037, group standard deviation of 0.069, MZ ICC = 0.46 (p = 0.003), and DZ ICC = −0.27 (n.s.). For FD_original_, within-subject standard deviations are 0.021 mm, group standard deviations are 0.061 mm, MZ ICC = 0.56 (p = 0) and DZ ICC = −0.03 (n.s.). For FD_filtered,4-TR_, within-subject standard deviation is 0.034 mm, group standard deviation is 0.078 mm, MZ ICC = 0.46 (p = 0.003), and DZ ICC = −0.12 (n.s.). All of these statistics conform to the pattern of respiratory and motion properties being most similar within-subject, next-most-similar within monozygotic twins, and then less similar within dizygotic twins (and unrelated subjects).

Prior studies have linked BMI to FD_original_ (Hodgson et al., 2017) and found no major influence of respiratory rate on that relationship. Here, beyond the BMI-FD_original_ link, we have described a link between two other measures: variance in respiratory measures like ENV, and mean values of the motion measure FD_filtered,4-TR_. As a final investigation of the link between BMI and respiration and motion, we examined discordance between twin pairs for BMI, standard deviation in ENV, and mean FD_filtered,4-TR_. There was no relationship of discordance of BMI to discordance of motion or of variance in ENV in either MZ or DZ twins (Figure S6). However, there was a strong relationship between discordance in variance in ENV and mean FD_filtered,4-TR_, r = 0.78 for MZ (p = 0) and r = 0.64 for DZ (p = 0). These are consistent with a causal “state” interpretation whereby the respiratory events indexed by variance in ENV cause increased punctate and gradual head motion.

## Discussion

The popularity of functional connectivity techniques, and the fact that head motion confers systematic artifacts to fMRI data while also correlating with variables of interest, has made the identification of head motion and its correlates an important topic in human neuroscience (Hodgson et al., 2017; Siegel et al., 2017; Zeng et al., 2014). The challenge of controlling for this artifact has been compounded by the emergence of fast-TR sequences: motion estimates in fMRI images obtained with sub-second sampling are qualitatively different than the older, slower motion estimates, both because constant respiratory motion no longer aliases to the extent it used to, and because larger, sustained motions get subdivided and lost amidst the constant respiratory motion. The different appearance of these estimates, and the detection in large datasets of strong relationships between BMI and motion, have led some investigators to abandon the use of realignment estimates as measures of motion due to a suspicion that the estimates are so contaminated by pseudomotion as to be untrustworthy (Glasser et al., 2018). Other groups have begun to apply respiratory filters to the data, agnostic of the extent to which they are suppressing motion or pseudomotion (Fair et al., 2018; Siegel et al., 2017).

Motivated by an apparent reduction in low-amplitude real motion in our subjects who wear head molds (Power et al., 2019), we examined fast-TR data for evidence of respiratory pseudomotion, and came to recognize five distinct kinds of respiratory motion: constant real motion, constant pseudomotion, real motion at deep breaths, real gradual motions modulated by breathing depth, and a low-frequency pseudomotion at deep breaths. We now discuss some implications of these findings.

### On the low-frequency pseudomotion

The fact that a low-frequency wave appears only in the phase encode direction and seemingly only at and just after deep breaths is interesting for several reasons. First, because it exhibits all the signatures of pseudomotion. Second, because it appears exclusively in the phase encode direction and does not “contaminate” other realignment parameters. Third, because it seems at times to have no instantaneous correlate in terms of the respiratory belt (the wave oscillates regardless of whether the belt measure is unchanging or changing monotonically); if it reflects a respiratory phenomenon it may have to do with intrathoracic dynamics of the lung not evident in abdominal circumference. Fourth, because it was not evident in our cohort of children but was present in all adult cohorts. And fifth, because it was the largest amplitude form of pseudomotion we detected. Future studies with more comprehensive physiological recordings may determine the basis of this phenomenon. One potential use of these waves would be to identify them in phase encode parameters as an index of deep inspiration in fMRI data lacking respiratory monitoring.

### On constant respiratory motions

Constant respiratory motion is evident in all directions of unprocessed images, and appears in all realignment parameters, most prominently in the superior-inferior axis and the pitch rotation, and also in whichever of the other directions is the phase encode direction. In unprocessed images, respiratory displacement in the vertical direction is present in all scans. In the literature examining pseudomotion, when the left-right direction is the frequency encode direction, virtually no respiratory motion is seen in it, whereas when the anterior-posterior direction is the frequency encode direction, respiratory motion is seen (Brosch et al., 2002; Durand et al., 2001; Raj et al., 2001). These results collectively indicate that true respiratory motion is primarily occurring in the vertical direction, less so in the anterior-posterior direction, and minimally in the left-right direction.

In this setting, left-right respiratory motion in the HCP data is likely almost entirely pseudomotion, which would be consistent with the very high proportion of oscillatory variance that can be explained by BMI alone (approximately 50-65% of the variance). However, if this in-plane motion behaves as the low-frequency pseudomotion identified above, this pseudomotion may not “bleed” into the other motion parameters. Thus, one advantage of a left-right phase encode acquisition is that pseudomotion is “trapped” in a direction with inherently little motion. By contrast, an anterior-posterior phase encode direction will mix real motion and pseudomotion in ways more difficult to disentangle.

Very importantly, pseudomotion became dissociated from real motion during times of obstructive apnea. Whereas central apnea abolished oscillations of all kinds, obstructive apnea eliminated phase encode oscillations but enhanced anterior-posterior and vertical oscillations. This result makes sense physiologically. In central apnea, there is no respiratory effort and thus no head nodding, and also no air flow and thus no pseudomotion. In obstructive apnea, there is no air flow and thus no pseudomotion, but the subject will increase inspiratory effort as they labor against the obstruction, truly moving the head. Potentially, the congruence or incongruence of phase encode and vertical oscillations could be used to classify apneas or hypopneas, or to index arousal state, since presumably these obstructive events occur as the subject relaxes or falls asleep (Duran et al., 2001). If true, then one would expect at the transition to sleep not only punctate hypnic jerks, but also potentially sustained, rhythmic motions, which can be large in amplitude.

The amplitude of pseudomotion will vary according to acquisition parameters and subject characteristics like BMI or breathing rate. We will therefore only state that in the HCP data, omitting the phase encode direction entirely from motion calculations yielded estimates with 67% of the original amplitude, correlated at r = 0.92 with the original estimates.

### On punctate or gradual respiratory motions

In the unprocessed images it appeared that, much as tidal breathing can move the head up and down, deep inspirations can move the head, and gradual changes in the depth of breathing can modulate head position in correspondingly gradual ways. We used respiratory measures like ENV and RV to gauge such events, and found a strong relationship of variance in these respiratory variables to motion. This relationship is not attributable to pseudomotion: it was absent to weak when constant respiration was part of motion estimates, and only emerged strongly once respiratory frequencies were suppressed. These results, in fast-TR data, resemble those obtained elsewhere by use in several slow-TR datasets (Power, 2017; Power et al., 2018). This kind of motion is modestly related to BMI (r ∼ 0.30-0.35), but the reasons for this association are unclear; BMI did not scale with variance in the respiratory variables above, leading us to speculate that motions of all kinds are transferred more forcefully to the head in heavier subjects due to the relative inflexibility of their chests and abdomens.

### On reformulated motion estimates

We examined several reformulations of motion, in part to understand the contribution of various phenomena to overall motion, and in part because visualization of motion has been challenging in fast-TR data compared to slow-TR data. By suppressing respiratory motion, and aggregating position changes over several seconds, we drew out motions that were apparent in position estimates but which had largely been “invisible” once temporally subdivided and blended into constant respiratory motions. Our particular reformulation of FD_filtered,4-TR_ is one of many possible and reasonable ways to represent motion. The 4-TR differential was chosen mainly to obtain an effective TR comparable to TRs we were accustomed to in the past, but also because 4 TRs is the time over which, in the past, we could readily detect systematic distance-dependent influences of motion, presumably in part due to spin history effects (Power et al., 2014). These modified motion estimates are visually helpful for understanding events during a scan – the motions are visible, they visually match position changes, they often correspond to changes in respiratory traces (deep breaths, modulations of breathing depth), and they often have corresponding DVARS spikes and dips.

Sampling over multiple TRs could potentially be tailored to a subject’s respiratory rate. For example, assuming sinusoidal motion, an “effective” TR near the full respiratory cycle time will yield minimal respiratory motions compared to any other fraction of the cycle time, a topic discussed in detail in (Fair et al., 2018). For the mean respiratory period of 3.33 seconds (0.3 Hz) in this report, the 2.88 second effective TR we used is approximately the typical cycle time.

### On the directionality of motion and respiratory effects

The respiratory records of the HCP data are from a single elastic belt about the abdomen, prompting us to make certain caveats about the information it provides. Respiratory efforts can occur nearly independently at both the abdomen and chest (Quan et al., 1997; Randerath et al., 2017). For this reason, in the respiratory literature, current state-of-the-art noninvasive respiratory monitoring is accomplished via inductive or light plethysmography, which involves dual belts or dual optical measurements of abdominal and thoracic excursions. More invasive ways to monitor respiration include monitoring air flow via a mask, and measuring end-tidal pCO_2_ by mask or nasal canula. Accompanying pulse oximetry is standard (the HCP pulse oximeter provides only a waveform, not blood oxygenation). And often electrocardiography and/or electroencephalography are performed, depending on if the study is during awake or sleeping periods. There is thus a substantial disparity in the kinds of records we possess in HCP data in comparison to the redundant and precise records obtained by investigators specifically studying respiratory phenomena.

Is it likely the HCP records are missing respiratory phenomena, or, conversely, that non-respiratory phenomena are appearing in the abdominal belts? In the respiratory literature, chest and abdominal traces are typically congruent even if amplitude is low in one location, with one major exception: incongruence or phase shifts between chest and abdomen records are a defining feature of obstruction. Our respiratory records show nearly constant respiratory efforts other than sometimes transient apneas after very deep breaths, which are normal phenomena; we therefore suspect that little information was fundamentally lost by not having a chest belt, other than a way to define obstruction (beyond the dissociations in head motions above). However, it is possible that body motion caused apparent changes in respiratory traces that might resemble respiratory phenomena. Movement of the body can strain the abdominal belt, causing brief changes to the trace that could conceivably appear as a respiratory pause or breath (and sometimes subjects do make glottal stops during motions involving abdominal contracture, but these will probably tend to be brief and without a significant impact on ventilation). Without a redundant source of respiratory information, like a chest excursion or nasal cannula measure, we have no way of definitively identifying such “spurious” respiratory events.

However, one way of disambiguating real respiratory phenomena from “apparent” respiratory phenomena would be to monitor cerebral blood flow. Ventilation reduces the concentration of CO_2_ in the bloodstream (pCO_2_), and pCO_2_ is a potent modulator of cerebral blood flow (Hall, 2016). Increases of pCO_2_ induced by breath holds or CO_2_-enriched gas inhalation can cause cerebral arterial flow increases of up to 50%; such changes are associated with large (several percent) BOLD signal changes throughout the brain (Bright et al., 2009; Kastrup et al., 1999; Poulin et al., 1996). Thus, one way of verifying that, for example, a deep breath is actually a deep breath and not a modulation of the respiratory belt by non-respiratory motion, would be to examine the fMRI timeseries for evidence of pan-brain signal changes reflecting alterations of cerebral blood flow.

Pan-brain signal changes associated with respiratory trace abnormalities are widespread in the HCP fMRI data. An example run from one subject is shown in Figure 14, and several examples of deep breaths in other subjects are shown in Figure 15. These images convey what is routinely seen in the HCP scans. By contrast, many major motions seem to have no effect on the respiratory belt (Figure S7 shows several examples). Though likely there are times when respiratory records are altered by abdominal motion apart from breathing, such times must be a minor portion of respiratory “events”, and are unlikely to have played any significant role in the findings of this paper. Relatedly, since we visually checked all physiology records for quality as a selection criterion for this paper, we probably excluded from our analysis many traces that would have had significant issues in this regard.

**Figure 14:**
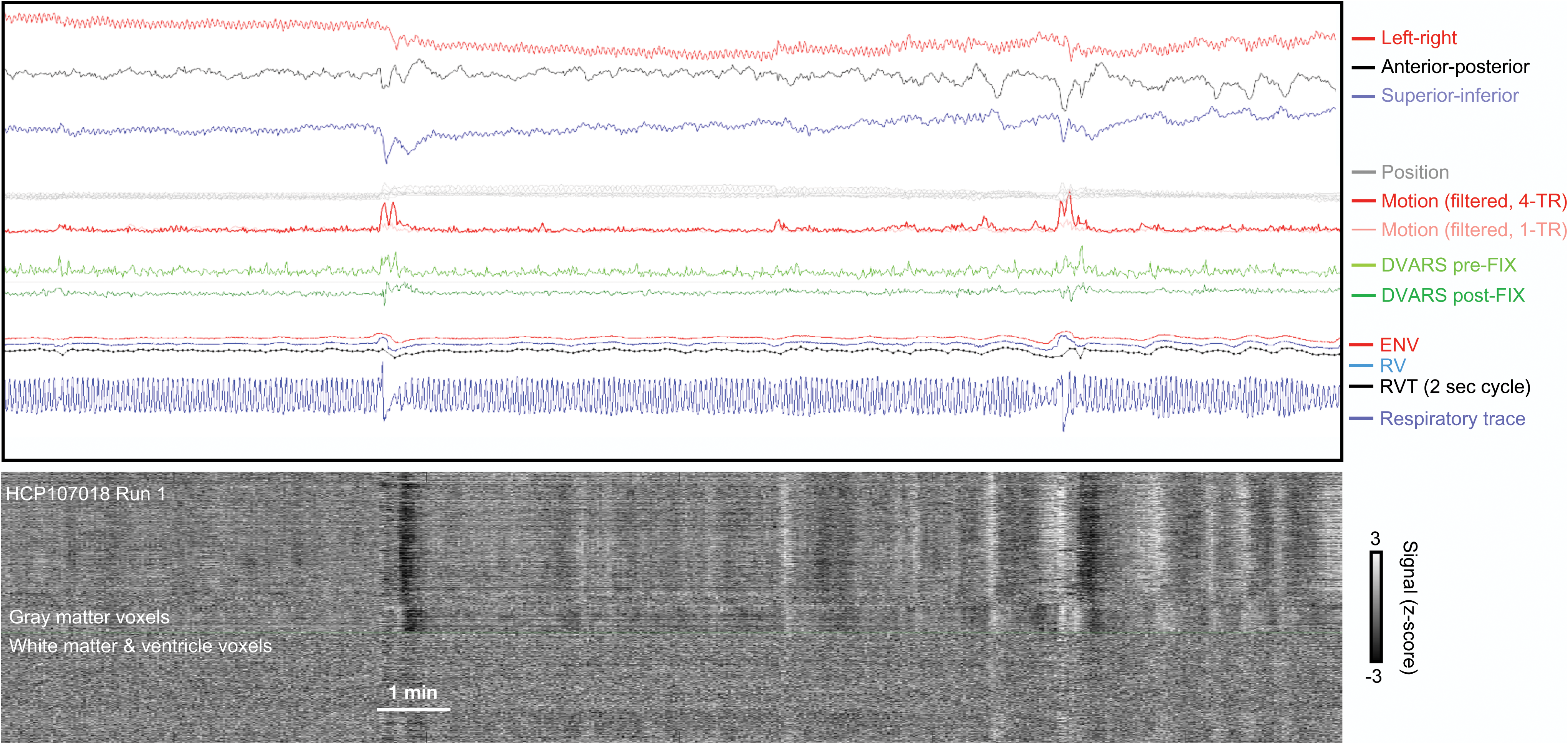
Concordance of respiration records and fMRI timeseries indicative of cerebral blood flow changes. Traces are as in prior figures, fMRI timeseries are from minimally processed image.

**Figure 15:**
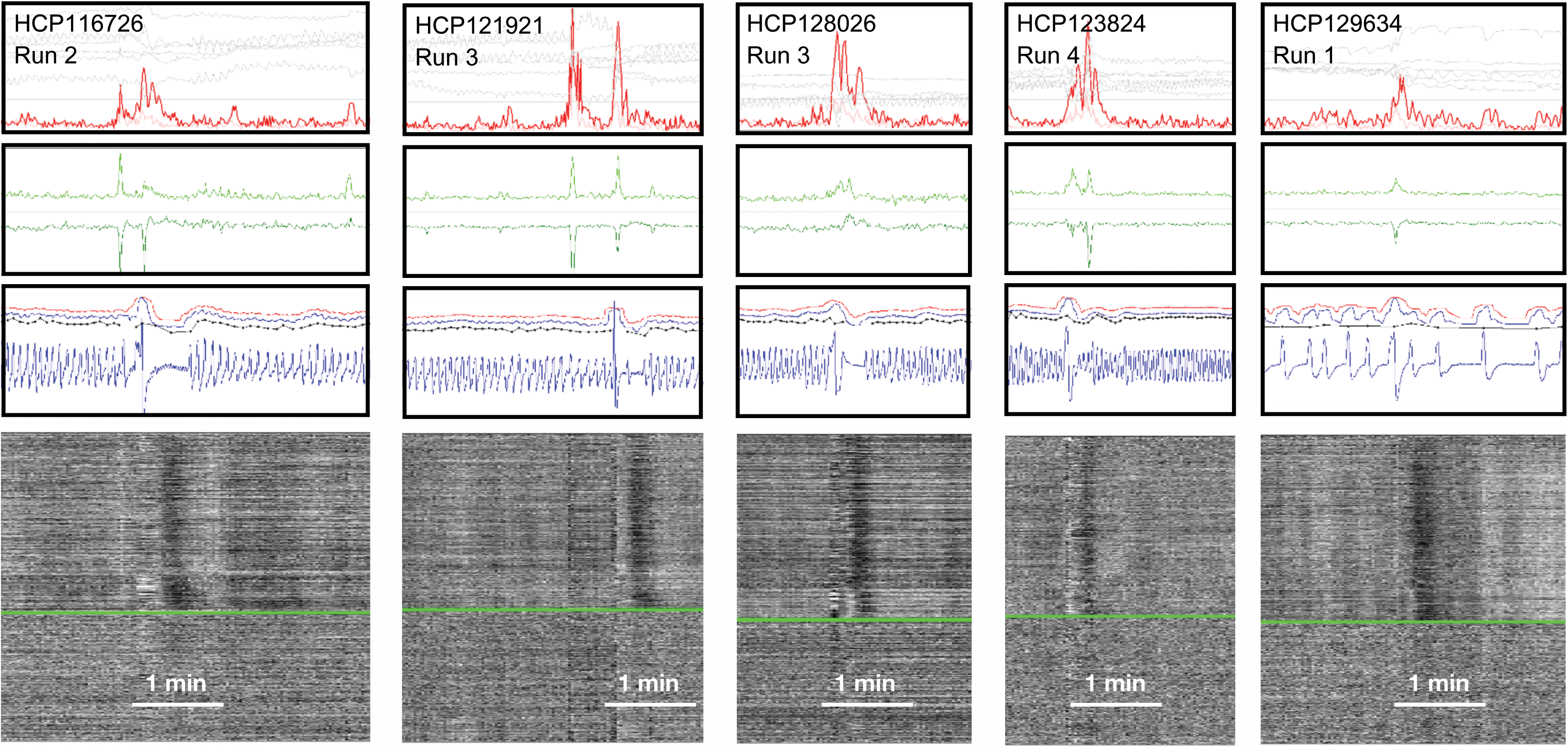
Concordance of respiration records and fMRI timeseries indicative of cerebral blood flow changes. Traces are as in prior figures, fMRI timeseries are from minimally processed images.

### On the fidelity of position and motion estimates in fast-TR data

We have uncovered evidence of pseudomotion in the HCP data, but this effect is limited to the phase encode direction. Though it is a substantial amount of the motion in the images (approximately a third of the magnitude), it is mainly oscillatory motion associated with respiration, and such oscillatory motion is also a true phenomenon in non-phase-encode directions. When the phase encode direction is not permitted to contribute to motion estimates, estimates correlate at r = 0.92 with the original estimates. We see little reason to mistrust the position estimates in HCP data, so long as the varied and distinct contributions of different kinds of motion and pseudomotion are kept in mind.

Our findings qualify the use of DVARS dips as an index of motion: DVARS dips are highly specific but not particularly sensitive indicators of motion. DVARS dips occur when FIX-ICA identifies structured abnormalities in the images sufficiently large to meet automated criteria, causing the denoising procedure to remove so much variance at those timepoints that those times exhibit unusually little variance thereafter. In our experience (and figures), essentially without exception, DVARS dips occurred at times of motion, and, in this sense, the dips are good indicators of motion. However, there are numerous instances of motion for which the dips are absent, mainly during smaller motions. Signal abnormalities during smaller motions are typically less marked than those during major abnormalities, and the FIX-ICA procedure was trained to mimic human decisions about components, which may explain why the denoising procedure readily identifies large motions that greatly disrupt signals in recognizable ways, but does not identify some smaller motions with less obvious signal disruptions. Note that DVARS spikes in minimally processed images very often accompany motions, indicating that signal disruptions are present, even if they are not recognized by FIX-ICA and converted to DVARS dips during denoising. A corollary of these observations is that DVARS-dip-associated findings are motion-related, but that non-DVARS-dip-associated findings are not necessarily not-motion-related.

### Revisiting the motivation for this paper

The investigations in this paper were prompted by reductions in motion and motion artifacts in fMRI scans obtained while using anatomy-specific head restraints (Power et al., 2019). In that report we contrasted our usual procedure of packing head coils with foam padding to restraining the head by filling the head coil with a customized Styrofoam insert molded to the subjects’ heads. In scans using head molds, absolute displacement and/or absolute motion were significantly reduced in all realignment parameters except the anterior-posterior axis, an effect we originally attributed to the head already being at rest on the occiput (we obtained modest but non-significant reductions in displacement and motion in the Y axis). Re-examination of those data leads us to suspect that the failure to substantially attenuate motion in the Y direction was probably due in part to pseudomotion. Without the mold, ongoing constant respiratory motion was prominent in the Z and Y directions, and with the molds the constant respiratory motion was visibly attenuated in the Z direction but not visibly changed in the Y direction (e.g., see Figure 1 or Figures S3-5 of (Power et al., 2019); Figure S8 reproduces Figure 1 with some modification). This observation prompts two comments. First, a developmental comment: though we did not see evidence of the low-frequency pseudomotion in our developmental Cornell 2 data (which was the mold “off” scans in only the younger subjects of (Power et al., 2019)), we do see the motion compatible with pseudomotion at the dominant respiratory rate in the data. And second, a comment on the utility of physical head restraints: to the extent that the residual motion in the Y axis with molds on was pseudomotion, we underestimated the reductions in “true” motion produced by the molds, which were a 60% reduction in large motions, and a 25% reduction in mean motion, when presuming all motion was “true” motion (Power et al., 2019).

### Broad considerations

Respiration can influence subject motion in multiple ways, and it influences fMRI signals prominently via cerebral blood flow modulations. It is hoped that resting state fMRI will one day have clinical relevance in psychiatry or neurology clinics. Preliminary studies indicate that resting state characteristics in extensively scanned subjects are stable and display unique features (Braga and Buckner, 2017; Gordon et al., 2017; Horien et al., 2019; Laumann et al., 2015). The same appears to be true for respiratory waveforms and presumably for the kinds of motion they produce: multiple within-subject parameters of respiration were most stable in individuals, next-most-stable in monozygotic twins, and less stable in dizygotic twins. Certain individuals had peculiar and idiosyncratic patterns in respiration (e.g., the deep breath and balloon envelope in Figure 4). In our data, the link between variance in summary respiratory properties (variance in ENV) and motion (FD_filtered,4-TR_) appeared independent of previously-described links of BMI and motion (FD_original_). Motion is a heritable property (Hodgson et al., 2017), and motion characteristics can classify individuals at above-chance levels in resting state studies (Horien et al., 2019). More studies will be needed to define the extent to which (heritable) respiratory phenomena contribute to such identifying features.

## Conclusion

We identified five kinds of respiratory motion in this paper, two of which are forms of pseudomotion, the other three of which are real motions. One form of pseudomotion is a low-frequency phenomenon not directly related to the respiratory rate. The other form occurs at the respiratory rate and is apparent in the phase encode direction of scans, but real respiratory motion is apparent in other directions at the same times. Motion estimates, once reformulated to accommodate the effects of various kinds of motion, yield productive information in fast-TR datasets that accords well with the information that they routinely delivered in slow-TR datasets in the past. The head is in constant motion in fast-TR images, but treatments of the motion estimates can permit investigators to “see beyond” the constant motion to identify meaningful events in scans.

## Acknowledgments

This work was supported by Simons Foundation Grant 528440 and a gift from the Mortimer D. Sackler, M.D. family. We thank Conor Liston, Henning Voss, and Steve Nelson for comments on the manuscript. We thank Kevin Tran and the NIH Helix/Biowulf staff for computing support.

## SUPPLEMENTAL

**Figure S1:**
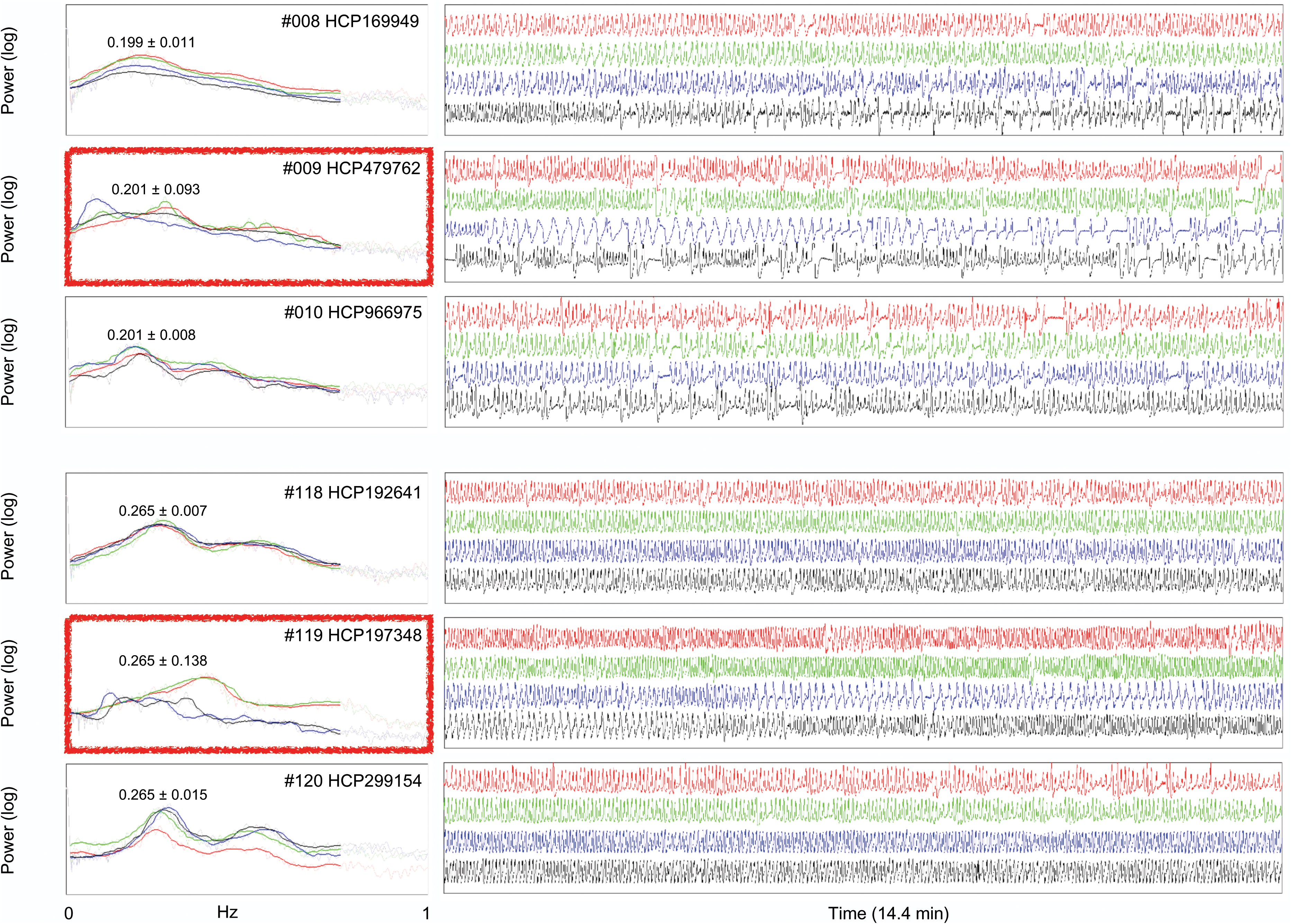
Examples of variable power spectra across runs in respiratory traces, after Figure 4. A minority of subjects have variable respiratory power spectra across runs (red boxes denote subjects with purple stars in Figure 4A with variable peak frequency). For reference, data from the neighboring ranked subjects are shown.

**Figure S2:**
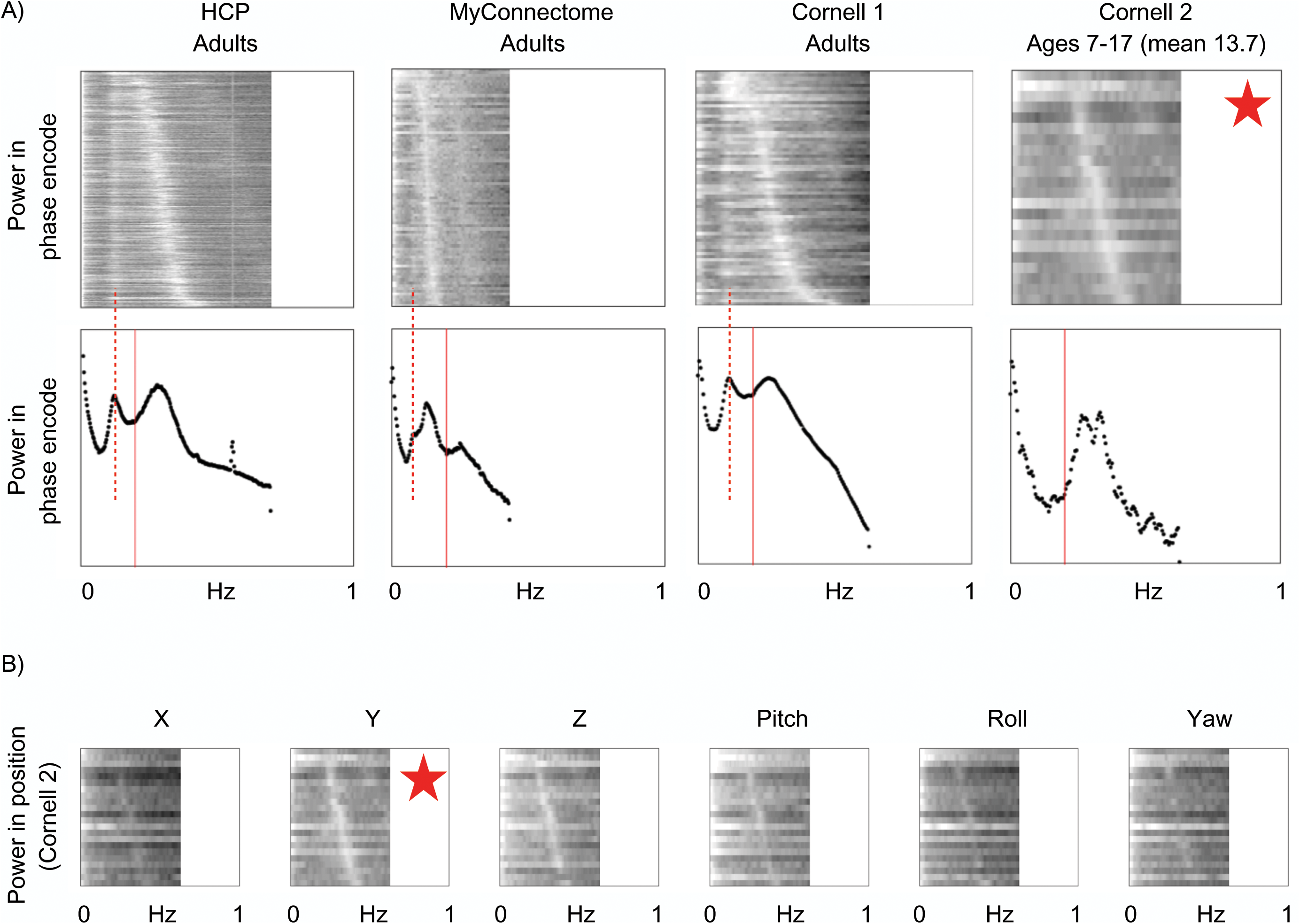
A) Position power spectra in the phase encode direction in adult versus developmental datasets. The Cornell 2 dataset is composed of 11 subjects ages 7-17, with 2 resting state runs lasting 4.8 min each per subject, TR = 800 ms, on the same scanner as the Cornell 1 dataset. In these children, there is no evident low-frequency peak. B) Position power spectra in all position estimates of the Cornell 2 dataset. Respiratory frequencies are represented in all position parameters, and no parameters show a visible low-frequency peak; the star denotes the phase-encode direction.

**Figure S3:**
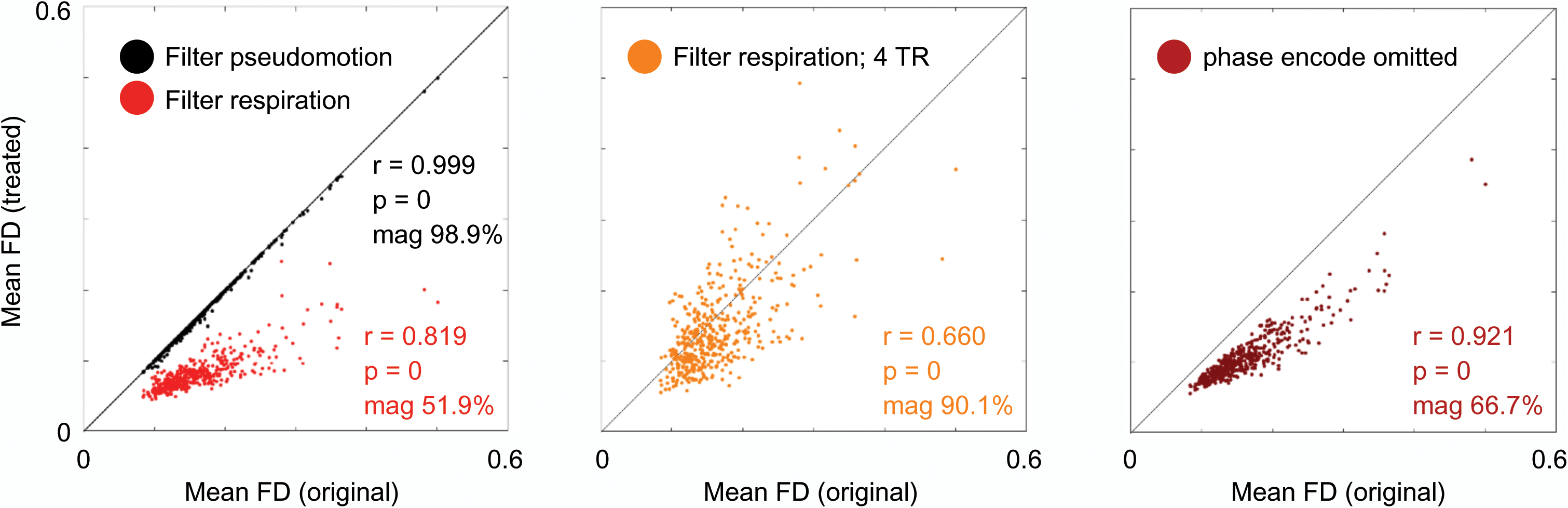
Summary motion during motion re-estimation. At right, mean FD calculated over the 4 runs of each subject defines the X-axis. Mean FD in the position estimates after filtering to remove either pseudomotion or the dominant respiratory frequencies is shown in black and red points, with summary statistics inset. At right, similar data are shown for the 4-TR FD estimates shown in Figure 11. In maroon are FD estimates produced without the X phase encode parameter

**Figure S4:**
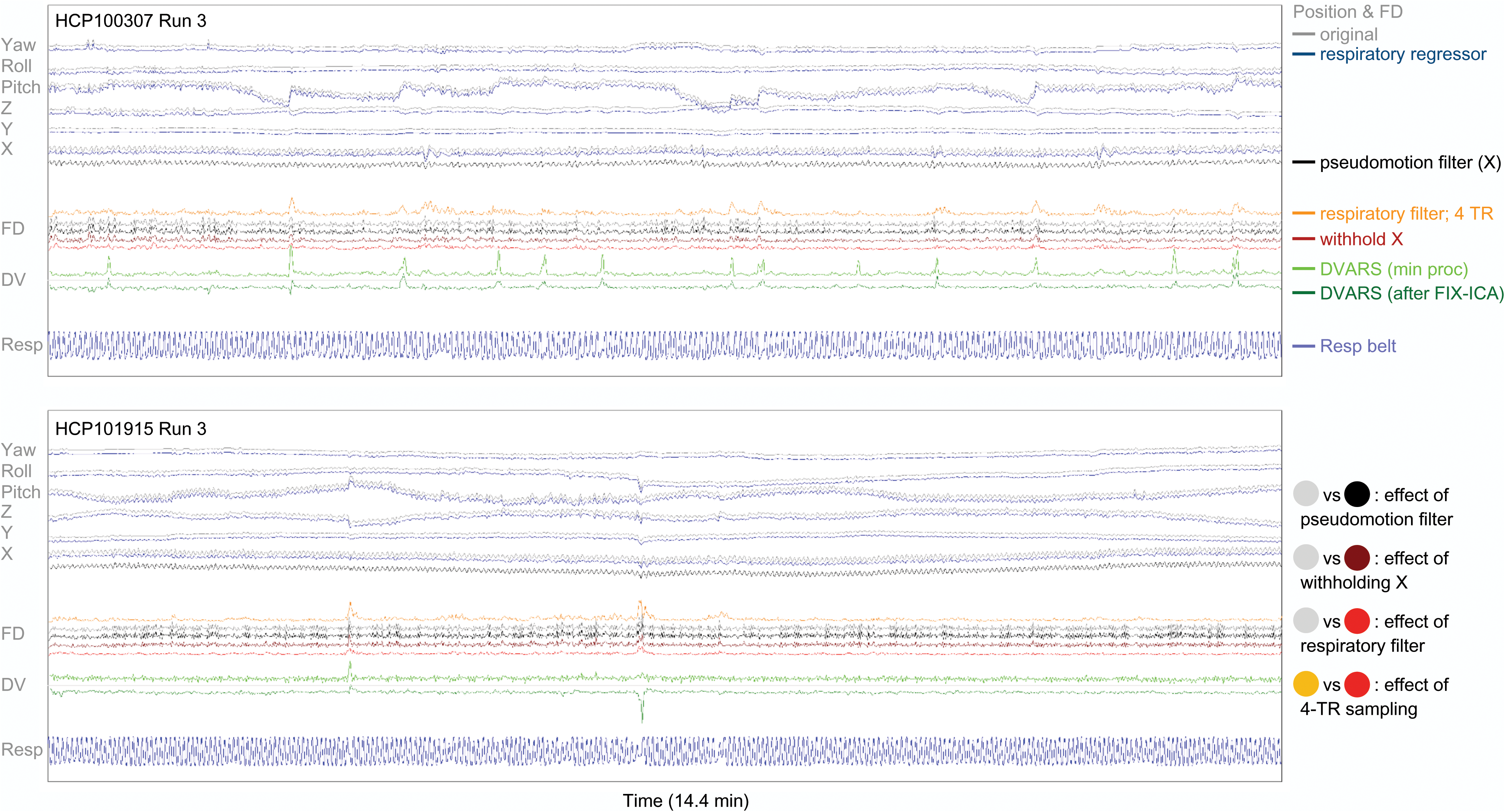
Figure in the style of Figure 10, but with blue position traces showing the effect of a respiratory nuisance regressor sampled from the respiratory belt instead of the red frequency-filtered position traces shown in Figure 10. The subject at top is the same subject shown in Figure 10, the bottom shows a different subject. The respiratory regressor attenuates respiratory oscillations best in the X parameter where the motion is most purely related to respiration (and in the top subject especially, revealing the distinct low-amplitude phase encode oscillations during and after some larger breaths). In subjects with marked excursions in the respiratory belt during deep breaths, the regressors tend to perform poorly.

**Figure S5:**
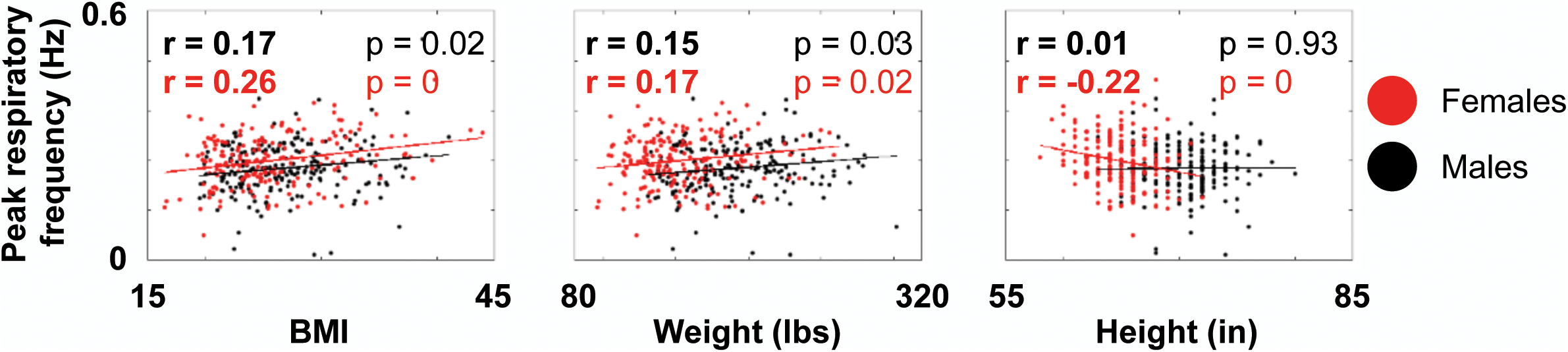
Interactions of body habitus with respiration rate.

**Figure S6:**
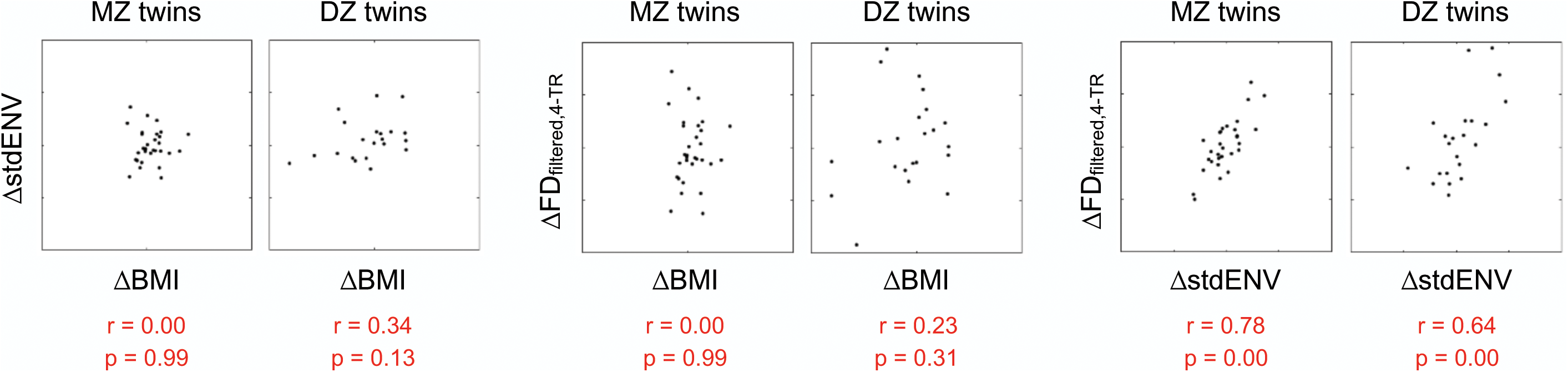
Discordance plots showing differences among twins on multiple variables. Discordance in BMI and ENV variance have no relationship to one another, discordance in BMI and mean FD_filtered,4-TR_ have no relationship to one another, but discordance in ENV variance and mean FD_filtered,4-TR_ have a strong relationship, supporting a causal link between respiratory phenomena and punctate/sustained motion.

**Figure S7:**
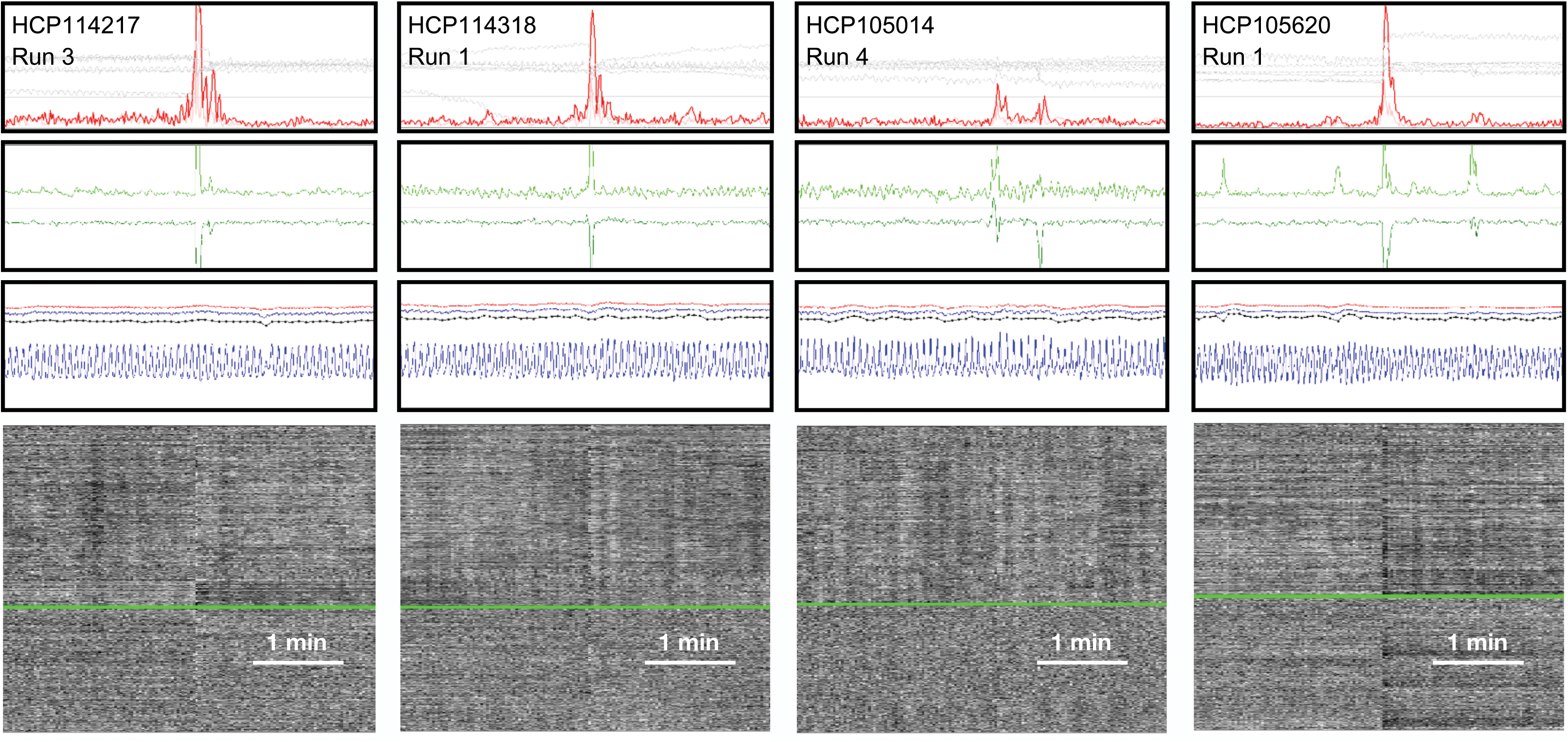
Instances of motion with no apparent relationship to respiration.

**Figure S8:**
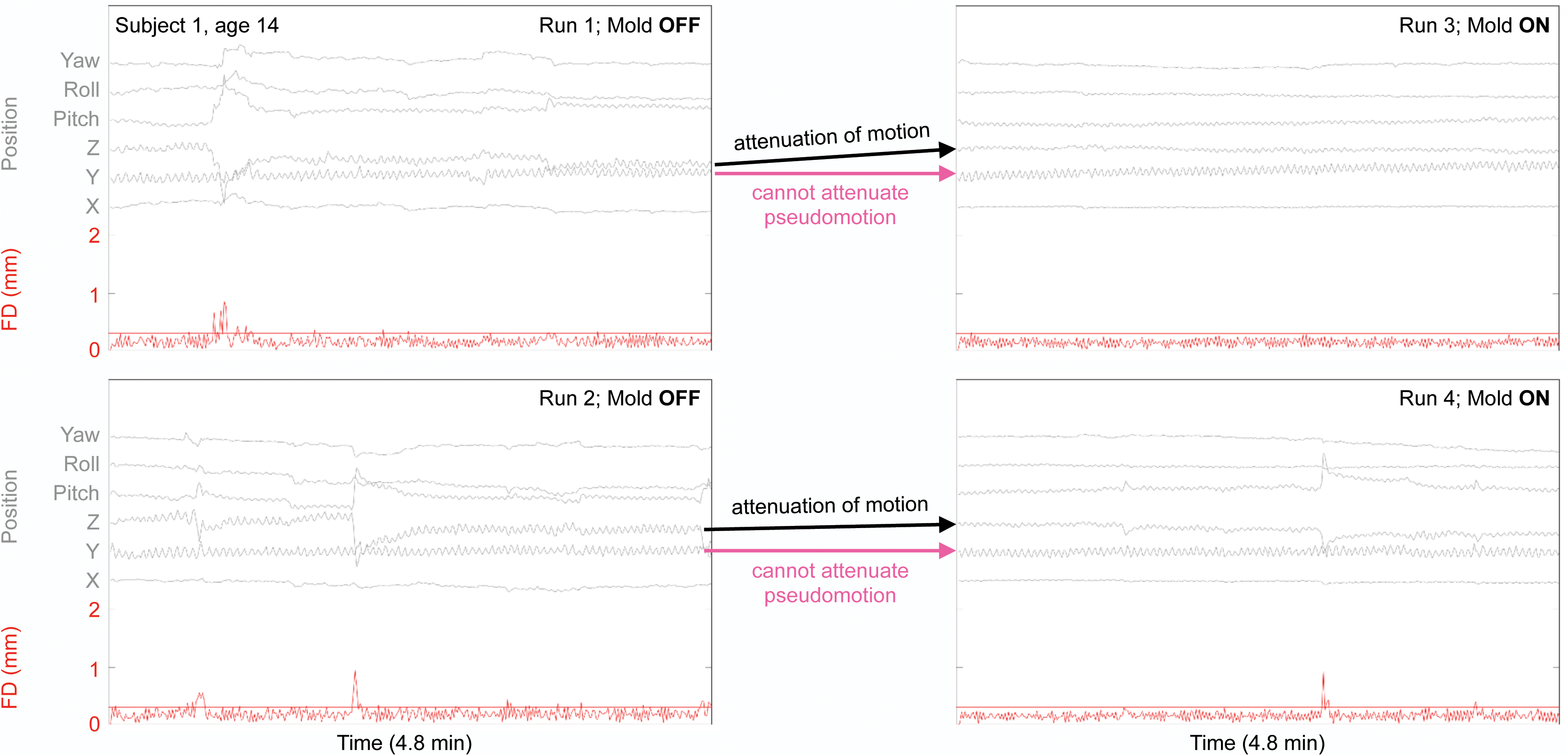
Position and motion estimates in the four runs of the first subject of {Power, 2019}. The horizontal red line is set at FD = 0.30 mm. Position estimates show respiratory motion prominent in the Y, Z, and pitch traces, as expected, most prominent with the mold OFF. With the mold ON, constant respiratory motion is visibly attenuated in the Z axis, but little changed in the Y axis, which is the phase encode direction in which pseudomotion would occur.

## Footnotes

1 https://www.jonathanpower.net/2018-glasser-comment.html

2 www.jonathanpower.net/2019-respiratory-motion.html

3 www.jonathanpower.net/2019-respiratory-motion.html

4 www.jonathanpower.net/2019-respiratory-motion.html

